# Hippocampal contributions to novel spatial learning are both age-related and age-invariant

**DOI:** 10.1101/2023.06.28.546918

**Authors:** Li Zheng, Zhiyao Gao, Stephanie Doner, Alexis Oyao, Martha Forloines, Matthew D. Grilli, Carol A. Barnes, Arne D. Ekstrom

## Abstract

Older adults show declines in spatial memory, although the extent of these alterations is not uniform across the healthy older population. Here, we investigate the stability of neural representations for the same and different spatial environments in a sample of younger and older adults using high-resolution functional magnetic resonance imaging (fMRI) of the medial temporal lobe. Older adults showed, on average, lower neural pattern similarity for retrieving the same environment and more variable neural patterns compared to young adults. We also found a positive association between spatial distance discrimination and the distinctiveness of neural patterns between environments. Our analyses suggested that one source for this association was the extent of informational connectivity to CA1 from other subfields, which was dependent on age, while another source was the fidelity of signals within CA1 itself, which was independent of age. Together, our findings suggest both age-dependent and independent neural contributions to spatial memory performance.

**Significance Statement:** Spatial memory declines with age, although some older adults show little cognitive decline, even into their 80s. One important lead for age-related changes comes from electrophysiological studies of older rats, who have less stable neural representations for a spatial environment, termed “place cells.” Using high-resolution fMRI targeting the human hippocampus, we demonstrate that older adults also have less stable neural representations within the same environment. Our results reveal, however, that there are both age and performance dependent differences driving these effects. While older adults, on average, showed alterations in information flow within the hippocampal circuit, better performing individuals showed more differentiated neural representations regardless of age. Together, these findings provide novel insight into the impact of age on spatial memory.

## Introduction

Numerous studies in rodents, nonhuman primates, and humans suggest that advanced age results in reduced spatial navigation performance and spatial memory declines compared to younger adults(1-4). Yet determining why such complex spatial memory difficulties(4) arise for many adults with advanced age has remained elusive. Single neuron recordings from rodents provide a potential lead, suggesting that some of these alterations may derive from differences in neural representations for spatial locations. For example, place cells in the CA1 subfield of the hippocampus show a reduction in the consistency of receptive field firing for the same environment(5, 6). This phenomenon, termed “remapping,” may relate to the known spatial memory difficulties of older rats and potentially humans, although this idea so far is untested in older adults. In the CA3 subfield of the rodent hippocampus, old rat hippocampal representations do not strongly distinguish between two different environments (i.e., de-differentiation)(7), suggesting a possible alteration in a computational mechanism termed “pattern separation.” This difficulty differentiating separate environments may relate more broadly to age-related network de-differentiation in human and nonhuman primate studies(8-12), although this, too, has not been studied across the lifespan.

Remembering different places involves formation of distinct neural representations for different environments and reinstatement of the same neural representation for the same environment. Therefore, if older adults show impaired spatial memory, one prediction is that reinstated activity patterns for the same environment might show lower fidelity between similar experiences than that of younger adults. On the other hand, the reinstated neural pattern for older adults in a different environment may be less differentiated compared to that of younger adults (Fig. 1D). Here, we used high-resolution functional magnetic resonance imaging (fMRI) targeting the hippocampus and other temporal lobe structures and employed multivariate pattern similarity (MPS) analyses. These analyses involve correlating the pattern of reinstated voxel activity within and between retrieved spatial environments. The hypothesis that decreases in older adult’s spatial memory might arise from alterations in both environment-specific codes and differentiation of neural signals between environments was therefore tested. Based on previous work(5-7), we focused on hippocampus subfields CA1, CA2 and CA3/dentate gyrus (CA2/3/DG) for these analyses; additionally, we tested parahippocampus cortex (PHC), entorhinal cortex (ERC), perirhinal cortex (PRC), and subiculum (SUB) as other critical medial temporal lobe subareas and inputs into CA2/3/DG and CA1.

**Fig 1.**
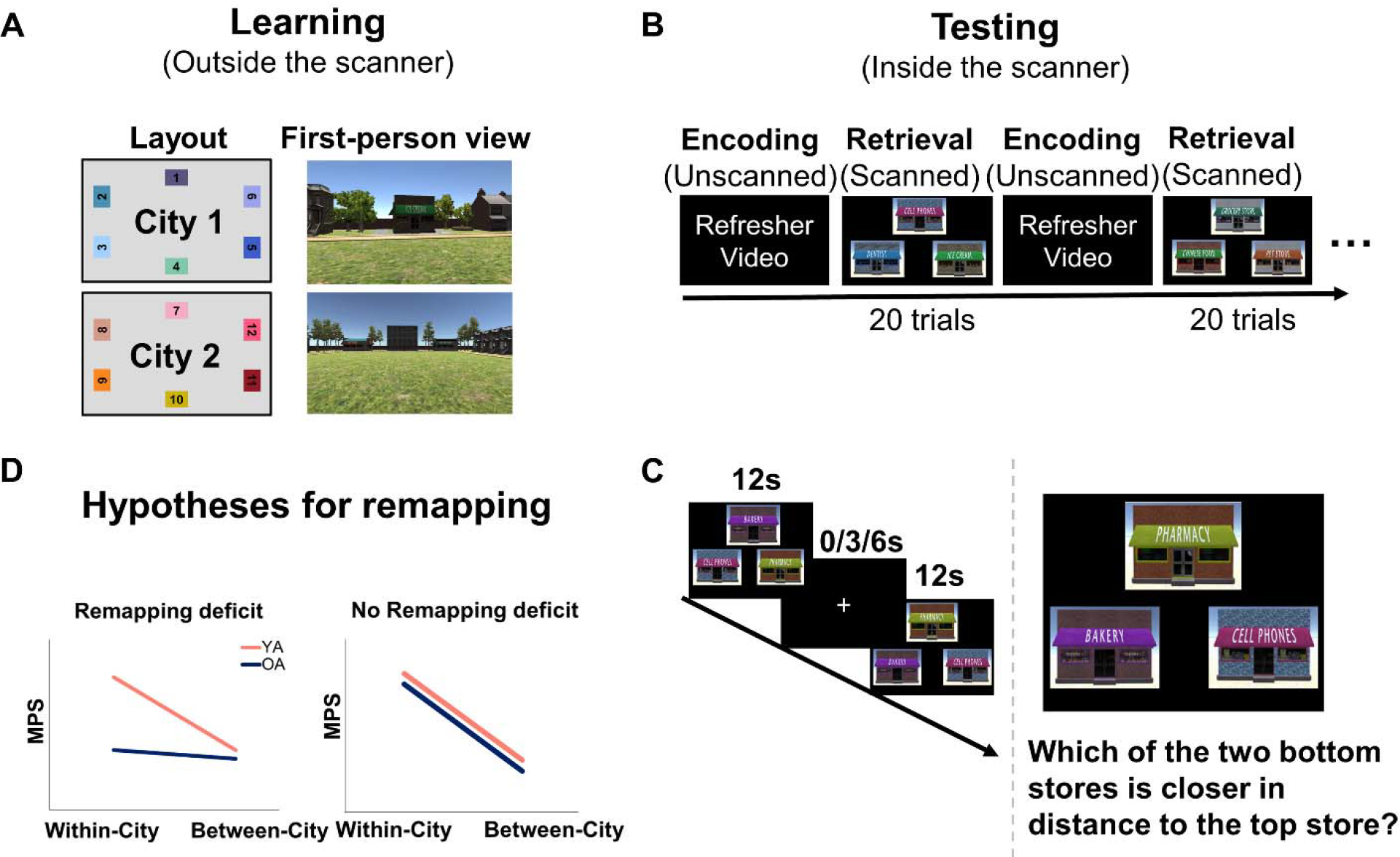
Schematic of the experimental design and hypotheses. A, Layouts for each environment are depicted from an overhead view (left) and first-person view (right). Each colored box represents a target store. All participants experienced the environments through the first-person view. B, Task structure in the MRI scanner. High-resolution fMRI scanning was only performed during retrieval and not encoding. The retrieval stage consisted of 6 runs of city-specific spatial distance discrimination, with three repeated runs per city. C, Participants were asked to perform a space distance discrimination task during the retrieval stage. D, Hypotheses, as laid out in the introduction: if older adults show impaired spatial memory, we should see alterations in the neural patterns for the same versus a different environment (i.e., de-differentiation: within-city MPS is comparable with between-city MPS in older adults) as well as less specific reactivation of the same environment (i.e., reduced within-city MPS compared to younger adults). YA: younger adults, OA: older adults.

In younger adults, more differentiated and specific neural patterns (as measured using MPS with high-resolution fMRI) correlate with more accurate retrieval of spatial distances within spatial environments(13, 14). The relationship between spatial memory performance and hippocampus subfield MPS, however, remains untested in older adults. Because older adults show individual differences in spatial memory with aging(15, 16), one possibility is that age-related differences in remapping may instead relate to poorer spatial memory, regardless of the age of the individual (“age-invariant”). For example, Schimanski et al. (2013) found that both older and younger rats with lower correlations in place cell representations in CA1 within the same environment also showed worse performance on the Morris watermaze(6). Alternatively, some age-related differences in remapping may also be “age-variant,” such that even when more accurately retrieving spatial memories for an environment, older adults continue to show differences in neural patterns. We investigated the mechanistic basis for these predicted effects by testing how differences in the input to CA1 might differ with age. We also measured how the degree of signal differentiation was related to CA1 neural remapping. Together, these measures allow us to determine both how changes in the inputs to specific hippocampus subfields, and the fidelity of signals within a subfield, relate more broadly to remapping.

## Results

### Older adults show lower and more variable multivariate pattern similarity (MPS) when retrieving the same environment compared to younger adults

Younger and older adults learned two different spatial environments to criteria (see Methods) and then retrieved the spatial distances of stores from these environments while undergoing high-resolution fMRI (Fig. 1). We performed multivariate pattern similarity (MPS), which assays how similar the voxel patterns are between different conditions (in this case, environments), on correct trials to test our hypotheses (see Methods). MPS was submitted to a linear mixed effects model with condition (within-city/between-city), ROI (PHC, ERC, PRC, SUB, CA1 and CA2/3/DG) and group (old/young). The model showed a significant condition by ROI by group interaction (*F*(5, 974483) = 8.232, p < 0.001, Table S1).

Consistent with our hypothesis, a simple main effects analysis revealed that within-city MPS was significantly greater than between-city MPS in CA1 for younger adults (*F*(1, 80.35) = 5.419, p = 0.022; Bayesian results: median = 0.0016, HDI = [0.0008, 0.0025], pd = 99.99%), but did not differ in older adults (*F*(1,130.93) = 0.447, p = 0.505; Bayesian results: median = - 0.00003, HDI = [-0.0012, 0.0012], pd = 51.91%, Fig. 2A). For younger adults, both within-city (t(41.6) = 4.587, p < 0.001; Bayesian results: median = 0.0045, HDI = [0.0029, 0.0061], pd = 100%) and between-city MPS (t(39.0) = 3.254, p = 0.002; Bayesian results: median = 0.0028, HDI = [0.0013, 0.0044], pd = 99.98%) were significantly higher than zero, whereas this was not observed among older adults (within-city MPS: t(52.5) = 0.720, p = 0.475; Bayesian results: median = 0.0004, HDI = [-0.0014, 0.0022], pd = 64.48%; between-city MPS: t(48.1) = 1.388, p = 0.171; Bayesian results: median = 0.0004, HDI = [-0.0014, 0.0022], pd = 66.40%). Furthermore, within-city MPS was significantly lower in older adults than in younger adults (*F*(1,47.33) = 6.076, p = 0.017; Bayesian results: median = 0.0042, HDI = [0.0017, 0.0067], pd = 99.95%, Fig. 2B), while there were no significant differences between the two age groups for between-city MPS (*F*(1,43.77) = 1.205, p = 0.278; Bayesian results: median = 0.0025, HDI = [0.0001, 0.0048], pd = 97.86%, Fig. 2C). We did not find this pattern in other hippocampus subfields or other medial temporal lobe areas (i.e., younger > older MPS within but not between city; see Supplementary Note 2 and Fig. S1 for the results from other medial temporal lobe areas). Together, the findings for CA1 suggest reduced consistency for the same environment during retrieval in older adults, in line with past findings from rodent studies(5, 6).

**Fig 2.**
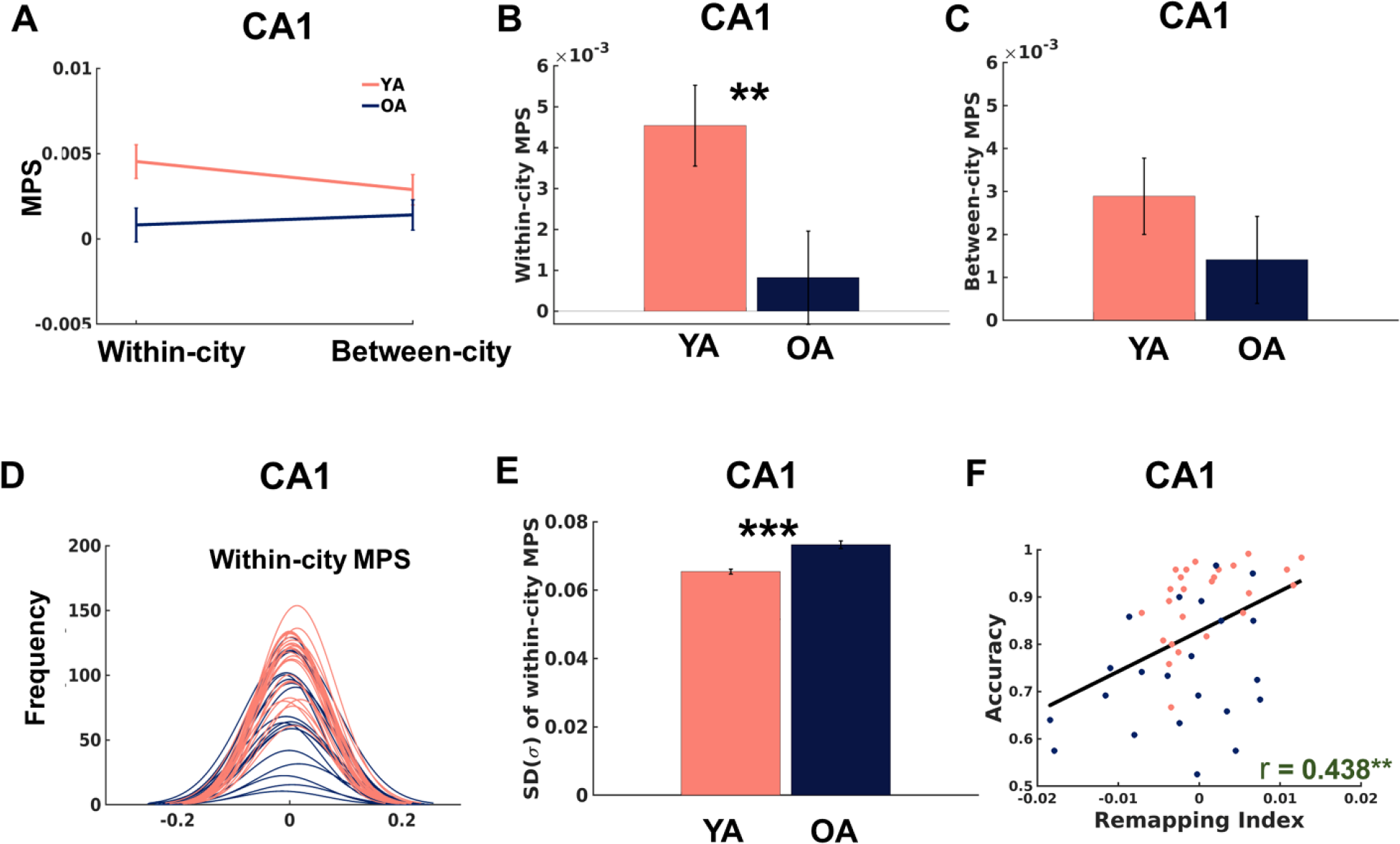
Differences in within and between city multivariate pattern similarity (MPS). A, Within-city MPS was significantly greater than between-city MPS in CA1 for younger adults but did not differ in the older adults. B, The within-city MPS in CA1 was significantly lower in older adults than in younger adults. C, The between-city MPS in CA1 was not significantly different in older and younger adults. D, The distribution of within-city MPS in the CA1 of older adults (dark blue) was wider than younger adults (red). E, Higher standard deviation of within-city MPS in CA1 for older compared to younger adults. F, The neural remapping index (within-city MPS minus between city MPS) in CA1 was positively correlated with spatial distance discrimination performance (“accuracy”). Each individual dot represents data from an individual participant, with red dots representing younger adults, while dark blue dots represent older adults. Error bars represent the standardized errors of the means. All data reflect n = 25 for YA, and n = 22 for OA independent participants. YA: younger adults, OA: older adults. **p < 0.01, ***p < 0.001.

Notably, spatial distance discrimination performance between older and younger adults differed, despite both groups beginning the experiment at criteria levels of spatial distance discrimination performance (see Methods), with older adults performing, on average, worse than younger adults (see Supplementary Note 1). Therefore, we used spatial distance discrimination performance and reaction time as covariates of no-interest. Importantly, we found that the within > between MPS interaction effect survived controlling for spatial distance discrimination performance and reaction time. In addition, we found that the age differences in MPS for within > between city could not be accounted by differences in univariate activation levels, activation variance, gender, ROI volume or greater shared perceptual similarity for within-city pairs (see Table S2 and Supplementary Note 2). In a control analysis, we re-examined all the aforementioned findings by including both correct and incorrect trials, which largely mirrored our findings with correct trials only (Supplementary Note 2).

Why should older adults, on average, show a lower correlation in voxel patterns for the same environment, even on correct trials (the focus of our analyses)? One possibility is that the neural representations are less stable for the same environment, as suggested in past studies in rodents(5, 6). To address this issue, we first examined the stability of neural representations by visualizing the distribution of within-city MPS on correct trials. The MPS of older adults showed a wider distribution of values than younger adults, indicating a broader spread of activation patterns within a city (Fig. 2D). Direct comparison of younger versus older adult within-city MPS revealed a larger standard deviation for within-city MPS in CA1 of older individuals (t(45) = 6.188, p < 0.001, Cohen’s D = 1.809; Fig. 2E). The Bayes factor calculated for the standard deviation for within-city MPS indicated extremely strong evidence against the null hypothesis (BF_10_ = 66530.829). This effect could not be accounted for by differences in performance when using spatial distance discrimination performance as a covariate in a linear regression model (R^2^_adjusted_ = 0.435, *F*(2, 44) = 18.734, p < 0.001). In control analyses, we re-examined the age-differences in the standard deviations of MPS when the number of trials were matched across the whole group and obtained similar results (Supplementary Note 2). Together, the findings above suggested that older adults showed a lower mean within-city MPS and higher within-city MPS variance, regardless of performance.

### Neural remapping and spatial distance discrimination are age-invariant

Next, we asked how age and performance related to the mean differences in correlations within and between cities, which can be quantified using the “neural remapping index.” A previous study from Kyle et al. (2015) showed that better performing younger adults showed a greater difference for within versus between city MPS (the “neural remapping index”). A higher score on this index indicated more differentiated environmental-specific neural representations (remapping) and a lower score indicated de-differentiation or less distinct network patterns. Previous work has suggested that older adults and nonhuman primates show greater de-differentiation for overlapping stimuli than do younger adults(8-12, 17). This may be particularly true for scenes, suggesting a parallel with navigation, although some studies have also supported the idea that these findings are age-invariant(18-20). In other words, better performing older adults could show comparable differentiation to younger adults, if the neural signals for the different environments are sufficiently differentiated.

Consistent with an age-invariant relationship between the neural remapping index and spatial distance discrimination, we found that the neural remapping index in CA1 was positively correlated with participant distance discrimination performance as measured by the percent of successfully retrieved distances of stores (r(45) = 0.437, p = 0.002, BF_10_ = 17.830, with strong evidence, Fig. 2F). This correlation was age-invariant (after controlling for age, gender, and CA1 volume, the correlation persisted, i.e., r_partial_(45) = 0.405, p = 0.006). We did not find any correlation between the neural remapping index in other MTL subregions (*ps* > 0.291, Fig. S5a; Supplementary Note 3). Together, these findings suggest that better performing participants – regardless of age – showed greater differences in the CA1 remapping index.

### Differences in the neural remapping index related to spatial distance discrimination are mediated by alterations in input to CA1

We then investigated potential mechanisms for the age-invariant correlations between the neural remapping index and spatial distance discrimination by considering how the neural signals in other MTL subregions might be contributing to this effect. Changes in connectivity are thought to underlie at least some age-related changes in hippocampus-mediated function(21). Reductions in the quality of input to CA1, for example, could result in less information available for local computations related to environment-specific codes. We quantified interregional informational connectivity using the values of the correlations of MPS between one subfield and another. Either higher or lower interregional connectivity was hypothesized as a plausible mechanism for reductions in input quality to CA1 (see Methods, Fig. 3A, B). We found that older adults showed increased interregional informational connectivity between PHC-ERC (t(45) = 2.467, Cohen’s D = 0.721, p_corrected_ = 0.03; BF_10_ = 3.172, with moderate evidence), PHC-CA1 (t(45) = 2.201, Cohen’s D = 0.645, p_corrected_ = 0.044; BF_10_ = 1.990, with anecdotal evidence), ERC-CA1 (t(45) = 4.249, Cohen’s D = 1.242, p_corrected_ = 0.001; BF_10_ = 207.124, with extremely strong evidence), ERC-CA2/3/DG (t(45) = 2.901, Cohen’s D = 0.848, p_corrected_ = 0.012; BF_10_ = 7.537, with moderate evidence), PRC-ERC (t(45) = 3.638, Cohen’s d = 1.063, p_corrected_ = 0.004; BF_10_ = 41.664, with very strong evidence), PRC-SUB (t(45) = 3.069, Cohen’s D = 0.897, p_corrected_ = 0.012; BF_10_ = 10.859, with strong evidence), PRC-CA1 (t(45) = 2.892, Cohen’s D = 0.845, p_corrected_ = 0.012; BF_10_ = 7.400, with moderate evidence), SUB-CA1 (t(45) = 3.018, Cohen’s D = 0.882, p_corrected_ = 0.012; BF_10_ = 9.716, with moderate evidence) and CA2/3/DG-CA1 (t(45) = 2.252, Cohen’s D = 0.658, p_corrected_ = 0.044; BF_10_ = 2.156, with anecdotal evidence, Fig. 3D). Here, we specifically focused on the connections between input areas (i.e., PHC, ERC, PRC, SUB and CA2/3/DG) to CA1 (Fig. 3C). We calculated the interregional informational connectivity by averaging across five selected connections: PHC-CA1, ERC-CA1, PRC-CA1, SUB-CA1 and CA2/3/DG-CA1 (see Methods and Fig. 3D). We found a negative correlation between the interregional informational connectivity and spatial distance discrimination (r(45) = -0.500, p < 0.001; BF_10_ = 90.288, with very strong evidence, Fig. 3E) as well as interregional informational connectivity and the neural remapping index (r(45) = -0.407, p = 0.005; BF_10_ = 9.092, with moderate evidence, Fig. S6e). This suggests that a higher correlation between patterns in different subfields related to worse spatial distance discrimination and a lower neural remapping index.

**Fig 3.**
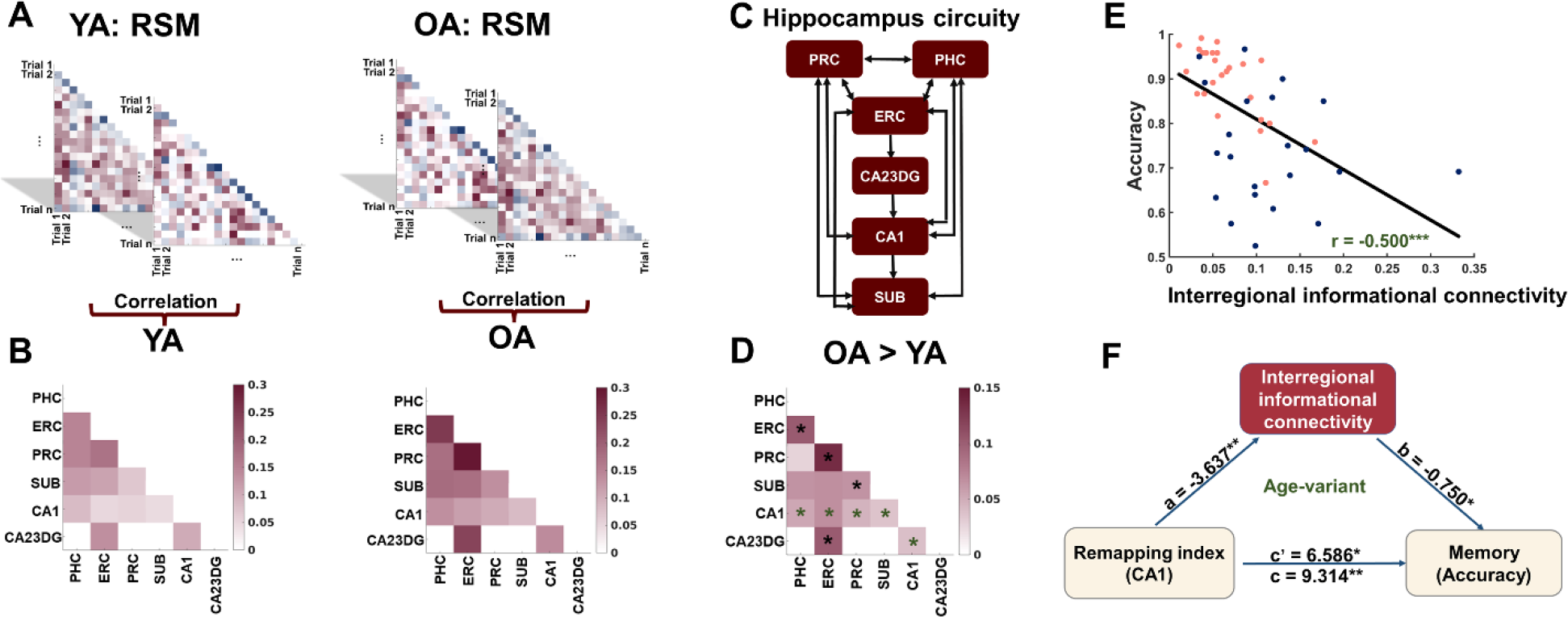
Differences in the neural remapping index related to spatial distance discrimination are mediated by alterations in input to CA1. A, Schematic of interregional informational connectivity analysis. For each ROI, we calculated the pairwise Pearson correlation among the activation patterns of trials to obtain a T-trial by T-trial representational similarity matrix (RSM) for each group (left: younger adults; right: older adults). B, A second-order representational similarity analysis was performed among the resulting RSMs for the 6 ROIs to obtain the informational connectivity matrix for each group (left: younger adults; right: older adults). C, Hippocampus circuitry (adapted from “Differential Connectivity of Perirhinal and Parahippocampal Cortices within Human Hippocampal Subregions Revealed by High-Resolution Functional Imaging” by Laura A. Libby, 2012.). D, Increased interregional informational connectivity in older adults from subfields with direct input to CA1 (the presence of asterisks (*) denotes statistical significance (p < 0.05, FDR corrected) for interregional informational connectivity in older adults compared to younger adults. Specifically, green asterisks highlight direct connections to CA1, while black asterisks indicate indirect connections with CA1. E, The interregional informational connectivity was negatively associated with spatial distance discrimination performance (“accuracy”). Each individual dot represents data from an individual participant, with red dots representing younger adults, while dark blue dots represent older adults. F, Interregional informational connectivity mediated the relationship between the neural remapping index (i.e., within-city minus between-city MPS) in CA1 and spatial distance discrimination performance (“memory accuracy”) after controlling for CA1 volume and gender, but this mediation effect was no longer significant after adding age as a covariate (i.e., age-variant). All data reflect n = 25 for YA, and n = 22 for OA independent participants. YA: younger adults (red), OA: older adults (dark blue). *p < 0.05, **p < 0.01, ***p < 0.001.

To attempt to better understand the relationship between interregional informational connectivity, the neural remapping index, and spatial distance discrimination, we performed a mediation analysis. The model indicated that interregional informational connectivity (Table S5) mediated the relationship between the neural remapping index in CA1 and spatial distance discrimination performance after controlling for CA1 volume and gender (Fig. 3F). Notably, when we entered age as a covariate, however, the mediation effect was no longer significant, suggesting that the interregional informational connectivity mediation effect was age-variant (Table S5). Similar results were found when using other connectivity patterns to CA1 (ERC-CA1, PRC-CA1, SUB-CA1, and CA2/3/DG-CA1); see Table S7, Supplementary Note 4, and Methods. We also examined the interregional informational connectivity between PRC-ERC, which has indirect connections with the CA1. The results suggest that PRC-ERC informational connectivity mediated the relationship between the neural remapping index in CA1 and spatial distance discrimination performance after controlling for CA1 volume and gender. This effect again did not survive after adding age as a covariate (Table S9), indicating that the interregional informational connectivity mediation effect was age-variant. We found similar results when considering the correlation only for the same environment, suggesting that this effect may have been driven by the stability of patterns for the same environment (Supplementary Note 4). Together, these findings suggest that differences in the input to CA1 contribute to age-variant differences in the neural remapping index.

### Correlations between the neural remapping index and spatial distance discrimination are mediated by reduced *CA1 neural* consistency

We estimated the dimensionality of neural signals by performing principal components analysis (PCA) to determine whether the fidelity of signals within hippocampus subfields also contributed to variance in spatial distance discrimination performance. We measured the dimensionality using the number of principal components (PCs) needed to explain 90% of the variance in voxel time course correlation patterns (Fig. 4A, see Methods). The results showed that the dimensionality of CA1 was significantly lower in older adults compared to younger adults (t(45) = 4.327, Cohen’s *D* = 1.265, p < 0.001; BF_10_ = 257.266, with extremely strong evidence, Fig. 4B). To ensure the robustness of this effect, we performed several control analyses. We selected the top 50 voxels based on each participant’s tSNR levels from CA1 (see Methods) to confirm that the age-differences we found were not influenced by differing numbers of voxels between individuals (Supplementary Note 5). We also performed the PCA analysis using all voxels within individual participant’s CA1 and matching the number of trials (Supplementary Note 5). These analyses again revealed lower dimensionality in older adults than in younger adults in hippocampus subfield CA1 (Supplementary Note 5).

**Fig 4.**
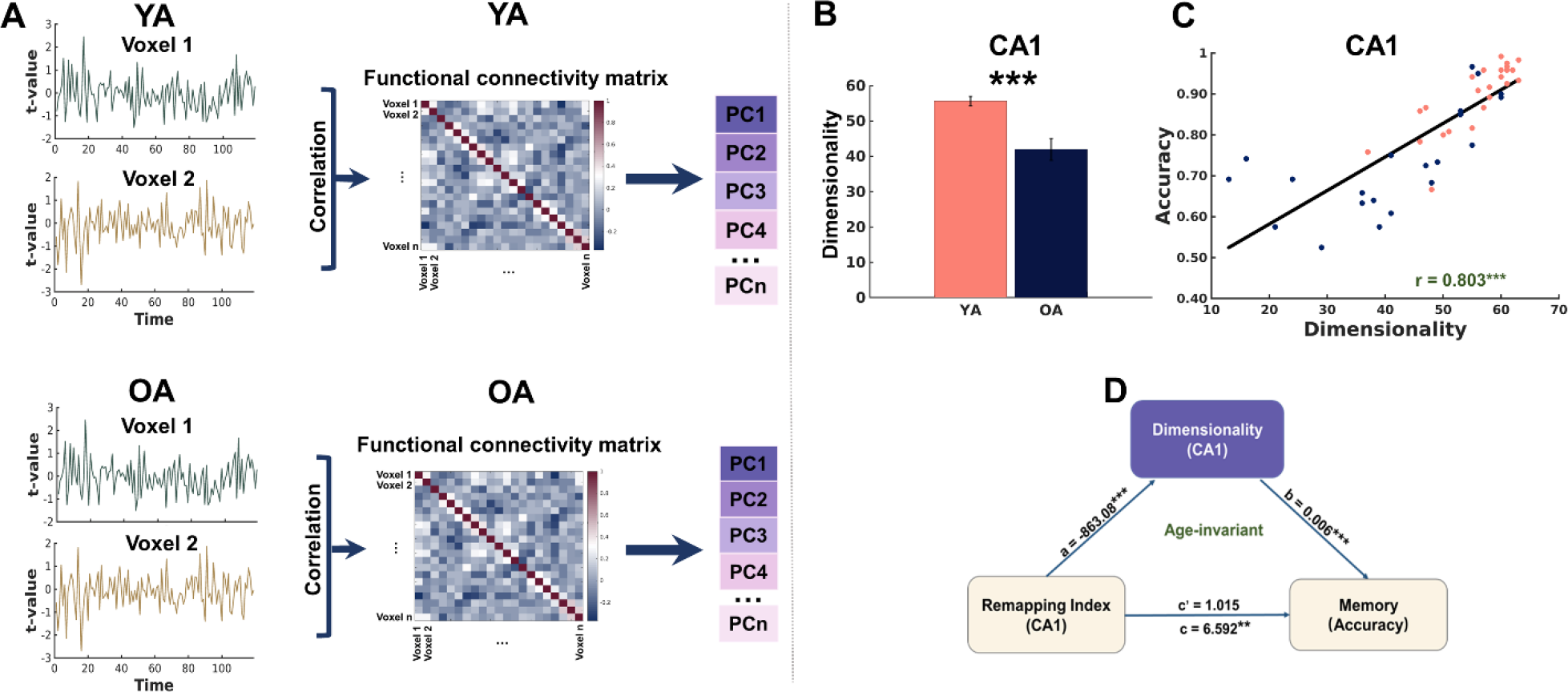
Correlations between the neural remapping index and spatial distance discrimination performance are mediated by reduced CA1 neural consistency. A, Schematic of principal component analysis (PCA): for each ROI, a series of t-statistics for each voxel of all correctly retrieved trials were extracted, and then used to calculate pairwise Pearson correlations, resulting in a functional connectivity matrix of N-voxel by N-voxel for each group (top: younger adults; bottom: older adults). PCA was then applied to the resulting N by N functional connectivity matrix to obtain PCs. B, Reduced CA1 neural consistency in older adults. C, The dimensionality of CA1 was positively associated with spatial distance discrimination performance (“accuracy”), even after controlling for age, gender and CA1 volume. Each individual dot represents data from an individual participant, with red dots representing younger adults, while dark blue dots represent older adults. D, The dimensionality of CA1 based on the PCA analysis fully mediated the effect of the neural remapping index (i.e., within minus between MPS) on spatial distance discrimination performance (“memory accuracy”), even after controlling for age, gender and CA1 volume. Error bars represent the standardized errors of the means. All data reflect n = 25 for YA, and n = 22 for OA independent participants. YA: younger adults, OA: older adults. PC: principal components. **p < 0.01, ***p < 0.001.

We found that CA1 dimensionality was positively associated with spatial distance discrimination (r(45) = 0.803, p < 0.001; BF_10_ = 911,300,000, with extremely strong evidence, Fig. 4C). Additionally, CA1 dimensionality was also positively correlated with the neural remapping index (r(45) = 0.538, p < 0.001; BF_10_ = 291.942, with extremely strong evidence, Fig. S7a). These effects were significant even after controlling for age, gender, and CA1 volume (spatial distance discrimination: r_partial_(45) = 0.722, p < 0.001; neural remapping index: r_partial_(45) = 0.495, p < 0.001), suggesting age-invariant effects. As discussed earlier, we also found a correlation between spatial distance discrimination and the neural remapping index in CA1 (Fig. 2F). Thus, we then ran an analysis to determine if the number of extracted principal components in CA1 mediates this effect. This analysis indicated that the dimensionality of CA1 mediated the correlation between the neural remapping index and spatial distance discrimination, even after controlling for CA1 volume and gender (Table S15).

The mediation effect remained significant when age was included as a covariate, suggesting that these effects were age-invariant (Fig. 4D, Table S15; see also Supplementary Note 5). These findings suggest that reduced fidelity of signals within CA1 mediated the relationship between the neural remapping index and participants’ memories for the distances of landmarks to each other in each city. While we did identify positive correlations between spatial distance discrimination and dimensionality in other MTL subregions (rs > 0.798, ps < 0.001, FDR corrected), we did not observe any significant correlations between the neural remapping index in other MTL subregions and dimensionality (ps > 0.08, FDR corrected), nor did we find any evidence of mediation effects in other MTL subregions (Supplementary Note 5). Therefore, increased neural differentiation between different cities was related to better spatial distance discrimination performance, based on the fidelity of signals within CA1 – regardless of age.

## Discussion

Our study focused on exploring the neural mechanisms underlying spatial memory decline in normative aging and how it might relate to well-documented changes in age-related hippocampus remapping in rodents. Because the neural representations within an environment should be more similar than those between environments, we subtracted the between city pattern similarities from the within city pattern similarities (generating the neural remapping index). We found that this index and spatial distance discrimination was positively correlated (Fig. 2F), reminiscent of the correlations found between CA1 remapping and spatial memory performance in rodents(5, 6). We found that one source of explained variance for this association was the extent of informational connectivity to CA1 from other subfields, which mediated the relationship of spatial distance discrimination performance and the neural remapping index yet was not present when controlling for age (Fig. 3F). A second source of variance in spatial distance discrimination performance was the fidelity of signals within CA1 itself, as measured by a principal components analysis, which partially accounted for worse performance and lower neural remapping index values (Fig. 4D). A mediation analysis that persisted even after controlling for age suggested this effect was age-invariant. Thus, spatial memory changes in older adults appear to occur for at least two reasons. Among these are changes in the quality of input to CA1, which differs based on age, and the other is related to the fidelity of signals within hippocampus CA1, which is independent of age.

Previous research using high resolution resting state fMRI has found that there are no significant differences in patterns of functional connectivity between younger and older adults within the hippocampus subfields(22). Our task-based functional connectivity analysis using the t-statistics time series method (Supplementary Note 6) also found no differences in connectivity patterns between the age groups. However, the interregional informational connectivity approach employed here (see Methods) allowed us to assess the extent to which two brain regions communicate similar or different information based on the similarity of voxel patterns between subfields(23). More similar voxel patterns between subfields that provide input to CA1 in older adults would result in reduced informational transfer to this subfield. This, in turn, would reduce the distinctiveness of representations for each environment in hippocampus subfield CA1. Specifically, the results suggest that the patterns of input to CA1 are important for spatial distance discrimination, and this effect varies with age.

Prior research points strongly to the idea that older rats show prominent differences in several electrophysiological measures. The place cells in CA1 of aged rats show less stable firing patterns between different exposures to the same environment, suggesting that the stability of place cell firing declines with age in older rats(5, 6). Older rodents also show reduced hippocampal plasticity in the form of faster decay of long-term potentiation than do younger rodents, which is correlated with faster forgetting of spatial information in the Barnes maze(24). In the present study, we provide the first evidence that the stability of neural activity patterns in CA1 for the same environment was reduced in older adults (Fig. 2E) and was related to worse spatial distance discrimination performance (Fig. S4). Although fMRI is an indirect measure of neural activity and cannot assay single cell activity, our approach allowed us to image hippocampus subfield activity at higher resolution (1.6 mm x 1.6 mm x 2 mm) than many studies, with our focus on distributed populations of voxels within hippocampus subfields, which may be more closely related to the activity of ensembles of cells in rodents and monkeys(25, 26). Given the finding that age-related differences in input to CA1 mediate the stability of these patterns and their relationship to spatial distance discrimination, our results are consistent with the idea that age-associated cognitive impairments arise, in part, from underlying neurobiological changes possibly related to the stability of neural ensembles and neural plasticity.

While an accurate representation of a single environment is critical for maintaining a stable memory over time, reliable discrimination between environments is also critical for navigation. Previous findings in young humans and rodents(13, 14, 27, 28) support a role for CA1 in differentiating between spatial environments. Such findings have also been observed in memory tasks in older adults, with worse episodic memory performance being related to reductions in neural differentiation(8, 10-12, 17-20). Older rats also show reduced differentiation of neural patterns in CA3 place cells compared to younger rats(7). The degree to which neural patterns differentiate between different environments can vary across individuals even in younger adults(14). Here, we found that better performing participants in both age groups showed more distinct neural patterns for different environments, consistent with the hypothesis that greater network differentiation is related to memory accuracy regardless of age.

The correlation between increased differentiation and better spatial distance discrimination performance was mediated by the dimensionality of voxel time courses in CA1. In our present study, we observed a decrease in the dimensionality of neural signals in CA1 among older adults compared to younger adults while engaging in a spatial memory retrieval task. These effects, however, were age-invariant, suggesting that better performing older adults had behavioral performance comparable to some younger adults. The reduction in neural signal dimensionality indicates the neural activity of voxels within CA1, which in turn may have a relationship to the differentiation of codes within that subfield. One possible interpretation of this finding is that less differentiated codes in subfields such as CA1 relate to worse spatial memory performance regardless of age. Unlike our earlier mediation results, which showed an age-variant relationship between differences in input to CA1 and spatial distance discrimination performance, the mediation by voxel dimensionality effect was not related to age. The origins of such within-subfield effects are unclear, but could relate to the content of available information within brain area, which could contribute to less accurate memory performance(29). Our data suggest that richer variability in neural activity within a subfield may support better differentiation of spatial representations and more accurate spatial retrieval. Our findings also suggest that alterations in input to hippocampus CA1 contribute to the well-documented spatial memory declines with age.

We note some important limitations in our study. The first is that older adults showed within and between city MPS that did not differ from zero, despite using correct trials. While the same effects were present when using correct and incorrect trials, the near zero MPS for the same and different cities in older adults suggest potentially that the pattern similarity analyses may show an exaggerated group difference. In other words, given that older adults, on average, performed well at the task, it is not clear why the pattern similarity results would not be different than zero. One possibility could be the greater variability present in older adult neural patterns for the same and different cities compared to younger adults (Figure 2E). Between-subject designs, such as that employed here with respect to age, involve some confounding factors, such as cohort effect and survivor effects (30). Also, the results of mediation analyses should be interpreted in light of the fact that they based on cross-sectional data, which limits their ability to reveal the causal structure of longitudinal change (31, 32). Another limitation is our modest sample size, which could have masked some smaller-effects sizes, for example, that older adult pattern similarity values might be above zero for the same environment with sufficient data. The challenge associated with between subject designs could be addressed in a future study using a longitudinal design involving the same older adults to examine how within and between city correlations might change with age. Finally, the relationship between task-related functional connectivity and underlying synaptic physiology remains unclear. As such, it is difficult to know how to interpret the finding that PHC/PRC functional connectivity mediated the relationship between neural remapping in CA1 and spatial distance discrimination performance because functional connectivity indicates both direct and indirect connections(33). We speculate, however, that input from PHC and PRC, who may have more of a direct role in relaying sensory information(34), may be more relevant to hippocampal processing than those further along in serial processing.

## Materials and Methods

### Participants

Twenty-eight younger adults and twenty-five older adults were recruited from the Tucson community and the study procedures were approved by the Institutional Review Board at the University of Arizona. Written informed consent was obtained from each participant before the experiment. Three participants were excluded due to technical issues. Two participants were excluded from analysis due to too few trials after motion correction. One participant was excluded due to voluntary withdrawal during the MRI scan. The final sample consisted of 25 younger adults (14 females; mean age: 23.96 years, range: 18-34 years) and 22 older adults (13 female; mean age: 72.68 years, range: 65-82 years).

All participants were right-handed and had normal or corrected vision. Based on self-report, all participants were screened to ensure they were in good health and had no neurological or psychiatric conditions. Older adults were also screened for dementia using a battery of neuropsychological tests, including long-term memory using the California Verbal Learning Test (CVLT), language using Boston Naming Test scores, speed/executive function using Trail-Making Test Parts A and B, and visuospatial abilities using Rey Complex Figure Test (RCFT). Participants were considered to have MCI if they scored 1 SD below the mean on two measures within one cognitive domain (i.e., memory, language, or speed/executive function), or 1 SD below the mean on each of the three cognitive domains. All the scores showed that all older adult performance was in normative range. In addition, the Santa Barbara Sense of Direction (SBSOD) survey(35) was adopted to measure participants’ self-reported spatial and navigational abilities (see Supplementary Note 1).

### Stimuli

We designed two different virtual environments using Unity3D (https://unity3d.com). Each virtual environment (i.e., City 1 and City 2) contained 6 different target stores located on the edge of the rectangular city (Fig. 1A, left). City 1 and City 2 shared the same basic layout, including the same ground, wall textures, and filler buildings but varied in terms of the identity of the 6 stores. Three first-person navigation videos were created for both cities (Fig. 1A, right).

Each video contained a unique, non-repeated navigation route.

### Experimental procedures

The experiment consisted of two sessions: a learning session outside the scanner and a retrieval session inside the scanner. During the learning session, participants were instructed to watch videos and to try to learn the store locations within each city. They were also informed there would be a spatial distance discrimination task to test whether they could successfully learn the relative distances between stores within each city. At the beginning of each round of the learning task, the participants were placed at the center of the city and viewed first-person navigation videos of travel from the center to each of the peripheral stores in a randomized order. Each learning task was followed by a shorter version (10 trials per run) of the memory retrieval task that they would later perform in the scanner (Fig. 1C). This learning-testing process repeated for each city in a randomized order for each participant until every participant met the criterion of 70% correct on at least one city before entering the scanner. Overall, 93.6% (N = 44) of the participants achieved 70% or higher correct responses for both cities while 6.3% (N = 3) of participants achieved 70% or higher correct responses for one city and 60% correct for the other city (see Table S17 for more details).

The fMRI retrieval session took place in the scanner, where participants completed six retrieval runs (three per city). Each run included 20 trials involving a single city and lasted 5 min and 2 s. Before the start of each retrieval run, participants viewed one of the city refresher videos (not scanned, Fig. 1B) which they had seen during the learning session to remind participants of which city they would be retrieving (scanned, Fig. 1B) next. The order of retrieval runs was randomized across participants. For each trial, participants saw three stores on the screen for 12□s, with one store on the top and two below (Fig. 1C, right). Participants were asked to compare which of the two bottom stores was closer to the upper reference store and indicate their choice by pressing the corresponding key on an MR-compatible button box. A “one” response indicated that the lower-left store was closer to the top store, and a “two” indicated the lower right store was closer to the top store. A trial lasted 12 seconds although participants were free to respond at any time. Between trials, presentation of stimuli was jittered using a central fixation cross with an intertrial interval of 0, 3, or 6 seconds.

After the spatial distance discrimination task, participants were asked to complete a control task involving vowel counting, which included two runs, each containing 20 trials. The structure of each trial in the vowel counting task was the same as in the spatial distance discrimination task.

Here, participants were asked to count the number of vowels (i.e., “A”, “E”, “I”, “O”, “U”; “Y” did not count) in each of the three store names and then select which of the two bottom stores had the closest number of vowels compared to the store on the top. Participants were asked to perform a vowel counting task as accurately and quickly as possible.

### MRI image data acquisition

Scanning was performed with a 32-Channel 3□T Siemens “Skyra” scanner located at the University of Arizona. Visual stimuli were projected onto a screen behind the scanner, which was made visible to the participant through a mirror attached to the head coil. Stimuli and responses were presented and recorded by PsychoPy (https://www.psychopy.org) on a Windows laptop. High-resolution anatomical images of the hippocampus and surrounding cortex were acquired with a T2-weighted turbo-spin echo (TSE) anatomical sequence: field of view (FOV)□=□200□mm□×□200□mm, matrix□=□448□× 448, repetition time (TR)□=□4200.0□ms, echo time (TE)□=□93.0□ms, flip angle□=□139 degree, slice thickness□=□1.9□mm, 28 slices, bandwidth□=□199□Hz/pixel. High-resolution functional images were acquired using a partial-brain echo planar imaging (EPI) sequence (interleaved acquisition, TR□=□3020□ms, TE□=□29□ms, flip angle□=□90 degree, FOV□=□192□mm, matrix□=□118□×□118, slice thickness□=□2□mm, slices□=□36, bandwidth = 1462 Hz/pixel), involving a voxel resolution of 1.6 × 1.6 × 2 mm. Sequences were acquired perpendicular to the long axis of the hippocampus. High-resolution structural images of whole brain were obtained using a 3D, T1-weighted MPRAGE (1□mm^3^ isotropic) sequence (FOV□=□256□mm, matrix□=□256□×□256, slice thickness□=□1□mm, TR□=□1800□ms, TE□=□2.26□ms, flip angle□=□8 degree, bandwidth□=□200□Hz/pixel).

### fMRI data preprocessing

Image preprocessing was performed by using FEAT (FMRI Expert Analysis Tool), version 6.00, implemented in FSL (part of the FSL package; http://www.fmrib.ox.ac.uk/fsl). The EPI images underwent motion-correction, temporal filtering (nonlinear high-pass filter with a 100□s cutoff), and slice-timing correction. Six motion parameters were added as confound variables to the model. Residual outlier timepoints were identified using FSL’s motion outlier detection program and integrated as additional confound variables in the first-level general linear model (GLM) analysis. No spatial smoothing was applied for single-trial estimation (see below). All functional images were linearly registered to the middle image of the first run and all analyses took place in native space.

### Single-trial response estimates

General linear models (GLMs) were performed separately to estimate the activation patterns for each of 120 retrieval trials. In this single-trial model, a Least Square–Separate (LS-S) approach was used, in which the trial of interest was modeled as one regressor, with all other trials modeled as a separate regressor. Specifically, each single-trial GLM included three regressors: (1) the trial during which a participant retrieved the spatial distance between stores; (2) all other remaining trials within the same run; (3) the remaining time period after participant made a judgement of all trials (i.e., 12s minus RT) while the stimulus stayed on the screen(36). Each event was modeled at the time of stimulus onset and convolved with a canonical hemodynamic response function (double gamma), whereas the fixation period was not coded and thus was treated as an implicit baseline. To control for the effects of head motion, six motion parameters were included in the GLM as covariates, as well as a regressor for each TR that was flagged as having greater framewise displacement (FD) than 0.5 mm during preprocessing. These generated t-statistics for each trial, with these t-statistics used for multivariate pattern similarity analyses (MPS), the principal components analysis, functional connectivity analysis, and interregional informational connectivity analysis to increase the reliability by normalizing for noise(37). The t-values were generated by dividing the beta value (regression fit) for each trial by the square root of the variance of residuals.

### Subfield demarcation and ROIs

Automatic hippocampus subfield segmentation software (ASHS) was used to segment the subregions of the MTL based on each participant’s high-resolution T2-weighted MRI image(38, 39). We used ASHS with the ASHS-Princeton-1.0.0-Young-Adult atlas in both younger and older participants. The MTL was segmented into CA1, CA2/3, DG, and subiculum (SUB), perirhinal cortex (PRC), entorhinal cortex (ERC), and parahippocampus cortex (PHC). We combined the CA2/3 and DG subfields as finer distinctions cannot be made at the acquired resolution(40). Each participant’s subfield segmentations were manually inspected to ensure accuracy of both segmentation protocols. Then, each subfield ROI was transformed into each participant’s native space using Advanced Normalization Tools (ANTs v2.3.5). Single-trial t-maps were then obtained within 6 ROIs (PHC, ERC, PRC, SUB, CA1 and CA2/3/DG) for each participant for further analysis.

For meaningful between group comparisons, it was necessary to register all brains to either the old or young adult atlas. This is because comparison between different groups requires analysis within the same atlas; otherwise, spurious effects can emerge due to atlas differences alone. To verify the reliability of our results, we also performed all the same analyses by using MTL ROIs that were segmented from ASHS with the ASHS-PMC-1.0.0-Old-Adult atlas in both younger and older participants. Note, results from the Young-Adult atlas are shown in the body of the manuscript and results from the Old-Adult atlas are shown in the Supplemental text.

### Multivariate pattern similarity analysis (MPS)

MPS was applied to measure the similarity of activation patterns by calculating the correlation between trials that were correctly retrieved in each hippocampus subfield. Following the approach of Power et al., we censored volumes with a framewise displacement > 0.5 mm and excluded any trials with any censored frames during the duration of the modeled GLM response(41). Specifically, for within-city MPS, pairwise Pearson correlation coefficients were calculated by correlating correct trials of the same city with each other. All between-city MPS analyses involved correlating correct trials between different cities. All MPS analyses involved correlating trials from different runs, thus avoiding temporal autocorrelations artificially inflating or biasing results. For the within-city calculation, we excluded trials that shared all three of the same store images (for example, triads “Store 1-Store 2-Store 3” and triads “Store 1-Store 3-Store 2”) to avoid identical images inflating correlations (See Supplementary Note 2). The resulting correlation coefficients were transformed into

Fisher’s z-scores. The standard deviation of MPS within each condition (within-city/between-city) was then calculated to depict the stability of neural representations of MPS between two age groups. The neural remapping index was defined to be the difference between within-city and between-city MPS, i.e., within-city minus between-city MPS.

### Interregional informational connectivity analysis

To examine interregional informational connectivity, we calculated the correlations between the voxel patterns for 6 MTL ROIs(23, 42). First, for each ROI, we calculated the pairwise Pearson correlation among the activation patterns of trials to obtain a T-trial by T-trial representational similarity matrix (RSM, Fig. 3A). In this analysis, only correctly retrieved trials were included and any trials with an fMRI volume that exceeded the framewise displacement threshold of 0.5 mm were removed. Note, to exclude the effect of intrinsic fluctuations on between-ROI representation similarity, only across-run pairs of the resulting RSM were selected to conduct the second-order representation similarity analysis between the 6 ROIs (Fig. 3B).

In order to calculate interregional informational connectivity, we performed the following steps: 1) we included only the 5 subfields that directly input into CA1 (i.e., PHC-CA1, ERC-CA1, PRC-CA1, SUB-CA1 and CA23DG-CA1); 2) We selected the relevant edges by applying a p-value threshold of 0.05 (i.e., p < 0.05, uncorrected) to the contrast of group differences. 3) we averaged interregional informational connectivity values to yield a single index for each participant. We calculated the interregional informational connectivity index for both the Young-Adult atlas and the Old-Adult atlas separately and identified unique subfields for each. We further used a relaxed p-value threshold of p = 0.1 to select common inputs for both atlases.

### Principal components analysis

Principal component analysis (PCA) was used to estimate the fidelity of voxel time courses within a subfield(43, 44). For each ROI, we extracted t-statistics for each correct trial, which reflected the fit of the hemodynamic response using the GLM described above. This generated a series of t-statistics for each voxel for all correctly retrieved trials. This N (voxels) by T (trials) matrix was then correlated with itself, generating an N-voxel by N-voxel matrix for that subfield (Fig. 4A). Note, similar to our MPS analysis, we also removed trials that included any volume that exceeded the framewise displacement threshold of 0.5 mm from the t-statistics series. Then, PCA was applied on this N by N functional connectivity matrix to obtain PCs using the MATLAB “pcacov” function (Fig. 4A). Finally, the dimensionality of a ROI was measured by calculating how many PCs were needed to explain 90% of variance, reflecting the potential fidelity of signals within that subfield.

For a given ROI, the maximum number of PCs is determined by the number of features (i.e., voxels). One potential issue in terms of the differences in dimensionality between two groups could be driven by the age-related differences in the number of MTL voxels. Therefore, we estimated the dimensionality by using a defined number of voxels in three different ways. First, for each MTL ROI, we obtained the voxel-wise tSNR(45) of the control task by calculating the mean of each voxel’s time series divided by its standard deviation. The voxels in each ROI could be ranked from high to low based on tSNR. After calculating each participant’s number of voxels for each ROI, we selected the smallest number of voxels as the defined number of voxels M for the whole group and matched the number of voxels by selecting the top M voxels based on the tSNR intensity. We repeated this procedure for each ROI, so each ROI had a matched number of voxels, M. Second, we arbitrarily selected the top 50 voxels based on tSNR for each ROI for two reasons 1) the dimensionality of all ROIs could be compared with each other since they all involved the same number of voxels and 2) compared to younger adults, the signal-to-noise ratio is relatively lower in older adults, and therefore it is possible that the age-related differences in dimensionality could be driven by the noisier voxels in older adults.

### Mediation analysis

Mediation analysis was performed using version 4.2 of the PROCESS macro(46) in SPSS (V28.0.1.1) to examine whether interregional informational connectivity between MTL ROIs mediated the relationship between the neural remapping index of CA1 and spatial distance discrimination performance. We performed similar analyses using principal components. Predictor (X) is the neural remapping index in CA1, spatial distance discrimination performance (Y) is the dependent variable, and interregional informational connectivity between MTL ROIs or the dimensionality of CA1 are mediators.

### Age-variant and age-invariant analysis

We systematically examined the presence of age-variant or age-invariant effects through a two-step approach. Initially, we conducted mediation analyses and/or correlation analyses without age as a covariate. In instances in which statistical significance was detected, a subsequent analysis was performed, incorporating age as a covariate. Results demonstrating sustained effects were categorized as age-invariant findings. Conversely, instances in which the observed effect was no longer significant upon the inclusion of age as a covariate were denoted as age-variant outcomes.

### Statistical Analysis

*Linear Mixed-Effects Regression*. Because the number of observations (i.e., pairs of MPS) is not balanced per condition (within-city/between-city) per participant, it is more appropriate to adopt linear mixed effects models to examine the remapping effect for younger and older adults(47).

Another advantage of applying a linear mixed effects model is that we can deal with confounding factors that could be contributing to the result on a trial-wise basis using covariates. For example, we can examine whether the differences for within-city MPS and between-city MPS were reliable after controlling for univariate activation levels and variance. Additionally, other possible confounding factors from individuals such as spatial distance discrimination performance, reaction time, and gender could also be included in the model as covariates to be regressed out.

Specifically, in R 4.2.2, we first constructed the maximal model with all possible random slopes by using the *lme4* package(48). Since the maximal model did not converge successfully, we simplified the model by removing random slopes gradually until it converged (BOBYQA controller, REML estimation). Therefore, the final model included random intercepts for each participant, random slopes for ROI (PHC, ERC, PRC, SUB, CA1 and CA2/3/DG), and/or random slopes for condition (within/between-city) for each participant. The significance of the model was evaluated using Satterthwaite approximation with the *lmerTest* package. Post-hoc tests based on the three way interactions were done using the *emmeans* package. We reported and plotted the estimated marginal means in the main test and figures.

*Bayesian analysis*. In addition to employing frequentist statistical linear mixed-effect regression, we also implemented Bayesian mixed-effects analysis to ensure the robustness of our results. We utilized the *brms* package(49) in R 4.2.2, employing 10 Markov chain Monte Carlo chains with 6000 iterations per chain, resulting in a total of 20,000 posterior iterations for coefficient estimation. Gaussian priors were employed to compute posterior medians, and we established 95% posterior credible intervals, also known as Highest Density Intervals (HDI), around the estimated effects. HDI that did not encompass zero were considered evidence of an effect. We also calculated the probability of direction (pd) using *bayestestR* package, which acts as an indicator for the presence of an effect and has strong correspondence with the frequentist p-value(50). Specifically, when the pd exceeded 97.5%, it indicated the presence of an effect equivalent to a two-sided p-value of 0.05. When the pd exceeded 99.5%, it signified significance at a p-value of 0.01, and when the pd exceeded 99.95%, it demonstrated significance at a p-value of 0.001. Additionally, we calculated Bayesian Factors using JASP (Version 0.16.4) for all t-tests and correlation analyses to further assess the strength of evidence for our findings. In all statistical tests, we applied a two-tailed test, and unless otherwise specified, we adopted false discovery rate (FDR) correction for multiple comparisons.

## Data, Materials, and Software Availability

All data is included in the article and/or supporting information and is available on the OSF (https://osf.io/zuqhj/).

## Acknowledgments

This research was supported by NIH/NIA grant R01 AG003376 to C.A.B and A.D.E.

