## Supplemental Tables and Figures for "Hippocampal contributions to novel spatial learning are both age-related and age-invariant"

**This PDF file includes:**

Supporting text
Figures S1 to S7
Tables S1 to S18
SI References

### Supporting Information Text

**Supplementary Note 1*:* Behavioral Results.** The accuracy of behavioral performance during spatial distance discrimination in the scanner was significantly lower for older adults than for younger adults (t(45) = 5.011, p < 0.001, Cohen's *D* = 1.465; $BF_{10}$ = 1826.288 with extreme evidence). Older adults were also slower to respond (t(45) = -2.419, p = 0.02, Cohen's *D* = -0.707; $BF_{10}$ = 2.901, with anecdotal evidence). The Santa Barbara Sense of Direction (SBSOD) was used to measure self-reported spatial ability, which showed a lower score in older adults than in younger adults (t(45) = -2.545, p = 0.014, Cohen's *D* = -0.744; $BF_{10}$ = 3.675, with moderate evidence). The SBSOD score, however, did not correlate with the neural remapping index on both atlases (the Young-Adult atlas: r(45) = -0.022, p = 0.883; $BF_{01}$ = 5.422, with moderate evidence for null result; the Old-Adult atlas: r(45) = 0.03, p = 0.837; $BF_{01}$ = 5.387, with moderate evidence for null result).

**Supplementary Note 2: Older adults show lower and more variable multivariate pattern similarity (MPS) when retrieving the same environment compared to younger adults**

*Older adults show lower MPS within the same environment in CA1 after controlling for covariates (results from the Young-Adult atlas)*: We used a mixed effects regression model to re-examine the remapping effect and age differences in MPS after controlling for univariate activation levels, activation variance, spatial distance discrimination performance, reaction time (RT), ROI volume, and gender when performing analysis using the Young-Adult atlas. The model showed a significant condition by ROI by group interaction (F(5, 974483) = 8.231, p < 0.001). Further, a simple main effect revealed that within-city MPS was significantly greater than between-city MPS in CA1 for younger adults (F(1, 82.2) = 5.396, p = 0.023; Bayesian results: median = 0.0019, HDI = [0.0008, 0.0025], pd = 100%), but did not differ in older adults (F(1, 134.5) = 0.811, p = 0.369; Bayesian results: median = -0.0001, HDI = [-0.0012, 0.0012], pd = 51.77%). For younger adults, both within-city (t(51.2) = 5.765, p < 0.001) and between-city MPS (t(48.2) = 4.377, p = 0.0001) were significantly higher than zero, whereas this was not observed among older adults (within-city MPS: t(69.1) = 0.531, p = 0.597; between-city MPS: t(63.5) = 1.986, p = 0.291). Furthermore, within-city MPS was significantly lower in older adults than in younger adults (F(1, 53.5) = 10.186, p = 0.002; Bayesian results: median = 0.0057, HDI = [0.0029, 0.0085], pd = 99.99%) suggesting lower reinstatement for the same environment during retrieval in older adults. However, no notable distinctions between the two age groups were found in terms of the between-city MPS (F(1, 48.7) = 3.525, p = 0.066; Bayesian results: median = 0.0040, HDI = [0.0012, 0.0068], pd = 99.73%).

*Older adults show lower MPS within the same environment* in CA1 (*results* from the *Old-Adult atlas):* To verify the reliability of our results and rule out the possibility that the results were driven by different segmentation protocols for boundaries of the MTL subregions, we repeated the same analysis as above using the Old-Adult atlas (See Methods). The mixed effects regression model also showed a significant condition by ROI by group interaction (F(5, 974474) = 7.412, p < 0.001, Table S3). For younger adults, both within-city (t(48.2) = 6.697, p < 0.001; Bayesian results: median = 0.0046, HDI = [0.0030, 0.0063], pd = 100%) and between-city MPS (t(41.7) = 5.081, p < 0.001; Bayesian results: median = 0.0032, HDI = [0.0017, 0.0048], pd = 100%) were significantly higher than zero, whereas this was not observed among older adults (within-city MPS: t(71.3) = 1.675, p = 0.098; Bayesian results: median = 0.0013, HDI = [-0.0006, 0.0032], pd = 91.61%; between-city MPS: t(58.5) = 1.471, p = 0.147; Bayesian results: median = 0.0002, HDI = [-0.0015, 0.0021], pd = 60.14%). Further simple effects revealed that within-city MPS was marginally significantly greater than between-city MPS in CA1 for younger adults (F(1,

86.25) = 3.337, p = 0.071; Bayesian results: median = 0.0014, HDI = [0.0005, 0.0023], pd = 99.81%), but not for the older adults (F(1, 142.52) = 0.095, p = 0.758; Bayesian results: median = 0.0011, HDI = [-0.0001, 0.0024], pd = 94.93%, Fig. S1c). Furthermore, older adults showed significantly lower within-city MPS than younger adults (F(1,60.4) = 8.673, p = 0.005; Bayesian results: median = 0.0033, HDI = [0.0008, 0.0058], pd = 99.52%) as well as lower between-city PS (F(1, 50.59) = 4.489, p = 0.039; Bayesian results: median = 0.0030, HDI = [0.0005, 0.0053], pd = 99.22%, Fig. S1d).

_Older adults show lower MPS within the same environment in CA1 after controlling for covariates (results from the Old-Adult atlas)_: Similarly, after controlling for univariate activation levels, activation variance, spatial distance discrimination performance, RT, ROI volume and gender, a significant condition by ROI by group interaction (F(5, 974518) = 7.829, p < 0.001, Table S4) was observed and further simple main effect revealed that within-city MPS was significantly greater than between-city MPS in CA1 for younger adults (F(1, 974510) = 5.635, p = 0.0176; Bayesian results: median = 0.0011, HDI = [0.0002, 0.0020], pd = 99.13%), but did not differ in older adults (F(1,974148) = 2.511, p = 0.113; Bayesian results: median = 0.0010, HDI = [-0.0003, 0.0023], pd = 93.73%). For younger adults, both within-city (t(101.2) = 7.285, p < 0.001) and between-city MPS (t(79.6) = 5.880, p < 0.001) were significantly higher than zero, whereas this was not observed among older adults (within-city MPS: t(110.1) = 0.644, p = 0.521; between-city MPS: t(85.4) = -0.604, p = 0.547). Meanwhile, older adults showed significantly lower within-city MPS than younger adults (F(1, 73.8) = 17.689, p < 0.001; Bayesian results: median = 0.0047, HDI = [0.0019, 0.0074], pd = 99.94%) as well as lower between-city MPS (F(1, 54.1) = 19.974, p < 0.001; Bayesian results: median = 0.0047, HDI = [0.0019, 0.0072], pd = 99.97%).

_Older adults show lower MPS within the same environment in CA1 when using all trials (results from the Young-Adult atlas):_ Besides, we recalculate the mixed-effects model for MPS by incorporating all trials (correct and incorrect), except those with an fMRI volume exceeding the framewise displacement threshold of 0.5 mm, to mitigate potential spurious correlations stemming from head motion. The results persisted with our initial results: a significant condition by ROI by group interaction (F(5, 1346576) = 10.185, p < 0.001) was found. Consistent with our hypothesis, a simple main effect analysis revealed that within-city MPS was significantly greater than between-city MPS in CA1 for younger adults (F(1, 49.52) = 4.206, p = 0.046), but did not differ in older adults (F(1, 59.36) = 1.953, p = 0.168). Furthermore, within-city MPS was significantly lower in older adults than in younger adults (F(1, 42.94) = 6.884, p = 0.012), while there were no significant differences between the two age groups for between-city MPS (F(1, 47.06) = 0.853, p = 0.360). Moreover, within the younger adult group, both within-city MPS (t(40.6) = 4.732, p < 0.001) and between-city MPS (t(44.5) = 3.796, p < 0.001) were significantly higher than zero, whereas within the older adult group, within-city MPS was not higher than zero (t(45) = 0.748, p = 0.458), while between-city MPS demonstrated a value exceeding zero (t(49.3) = 2.191, p = 0.033).

_Older adults show lower MPS within the same environment in CA1 when using all trials (results from the Old-Adult atlas):_ Besides, we recalculate the mixed-effects model for MPS by incorporating all trials (correct and incorrect), except those with an fMRI volume exceeding the framewise displacement threshold of 0.5 mm, to mitigate potential spurious correlations stemming from head motion. The results persisted with our initial results: a significant condition by ROI by group interaction (F(5, 1346576) = 10.185, p < 0.001) was found. Consistent with our hypothesis, a simple main effect analysis revealed that within-city MPS was significantly greater than between-city MPS in CA1 for younger adults (F(1, 45.07) = 3.459, p = 0.069), but did not differ in older adults (F(1,54.17) = 0.646, p = 0.425). Furthermore, within-city MPS was significantly lower in older adults than in younger adults (F(1,38.11) = 11.393, p = 0.002), while there were no significant differences between the two age groups for between-city MPS ($F(1,50.59) = 3.612$, $p = 0.063$). Moreover, within the younger adult group, both within-city MPS ($t(35.4) = 6.135$, $p < 0.001$) and between-city MPS ($t(47) = 5.356$, $p < 0.001$) were significantly higher than zero, whereas within the older adult group, within-city MPS was not higher than zero ($t(40.5) = 0.965$, $p = 0.340$), while between-city MPS demonstrated a value exceeding zero ($t(53.7) = 2.228$, $p = 0.030$).

_Older adults show lower MPS within the same environment in CA1 when using more stringent behavioral criterion for the training stage (results from both Young and Old-Adult atlases):_ We conducted a new mixed-effects model analysis to address the question that whether raising the behavioral criterion would lead to different results by excluding five older adult participants who did not meet a more stringent behavioral criterion (i.e., accuracy = 75%, see Table S17) for the training stage. The model showed a significant condition by ROI by group interaction (Young-Adult atlas: $F(5, 913477) = 7.934$, $p < 0.001$; Old-Adult atlas: $F(5,913466) = 8.844$, $p < 0.001$). Consistent with our hypothesis, a simple main effect analysis revealed that within-city MPS was significantly greater than between-city MPS in CA1 for younger adults (Young-Adult atlas $F(1, 68.97) = 5.327$, $p = 0.024$; Old-Adult atlas: $F(1,81.45) = 3.590$, $p = 0.062$), but did not differ in older adults (Young-Adult atlas: $F(1,106.88) = 0.037$, $p = 0.848$; Old-Adult atlas: $F(1,130.75) = 0.021$, $p = 0.885$). Furthermore, within-city MPS was significantly lower in older adults than in younger adults (Young-Adult atlas: $F(1,42.8) = 5.448$, $p = 0.024$; Old-Adult atlas: $F(1, 57.91) = 9.547$, $p = 0.003$), while there were no significant differences between the two age groups for between-city MPS (Young-Adult atlas: $F(1,39.21) = 1.960$, $p = 0.169$; Old-Adult atlas: $F(1, 46.65) = 3.533$, $p = 0.664$). All the results persisted after controlling for univariate activation levels, activation variance, spatial distance discrimination performance, reaction time (RT), ROI volume and gender as covariates: The model showed a significant condition by ROI by group interaction (Young-Adult atlas: $F(5, 913475) = 7.918$, $p < 0.001$; Old-Adult atlas: $F(5,913467) = 8.796$, $p < 0.001$).

_MPS results in other MTL ROIs after controlling for covariates (results from both atlases):_ We used a mixed effects regression model to re-examine the remapping effect and age differences in MPS after controlling for univariate activation levels, activation variance, spatial distance discrimination performance, reaction time (RT), ROI volume, and gender when performing analysis. On both the Young and the Old-Adult atlas, we observed higher within-city MPS than between-city MPS in PRC in older adults (the Young-Adult atlas: $F(1, 134.50) = 5.698$, $p = 0.018$; the Old-Adult atlas: $F(1, 974479.2) = 14.660$, $p < 0.001$, Fig. S1a and S1c) but not in younger adults ($p = 0.585$). However, we did not find any differences between the two groups in terms of both within-city MPS and between-city MPS (*ps* > 0.392, Fig. S1b and d). And we also did not find any relationship between the neural remapping index of PRC and spatial distance discrimination performance across the whole group (*ps* > 0.439, Fig. S5). Therefore, in the following analysis, we specifically focused on CA1.

_Control analysis for perceptual similarity of within-city pairs (result from both atlases):_ In order to exclude the possibility that the remapping effect was driven by shared perceptual information in the within-city condition compared to the between-city condition, we divided the within-city pairs into four categories: no shared store pairs ( i.e., no store that was overlapping between the trials used to calculate the within-city MPS, for example, triads “Store 1-Store 2-Store 3” and triads “Store 4-Store 5-Store 6”; one shared store pairs (i.e., only one store that was overlapping between the trials, for example, triads “Store 1-Store 2-Store 3” and triads “Store 4-Store 1-Store 6”); two shared store pairs (i.e., two stores that were overlapping between the pairs (for example, triads “Store 1-Store 2-Store 3” and triads “Store 4-Store 1-Store 2”); three shared store pairs (i.e., all three stores that were overlapping between the pairs (for example, triads "Store1-Store 2-Store 3" and triads "Store 1-Store 3-Store 2"). The more stores shared between trials, the higher the shared perceptual information, which could lead to higher pattern similarity. We then used a linear mixed effects model with condition (none-shared store pairs / one-shared store pairs / two-shared store pairs / three-shared store pairs), ROI (PHC, ERC, PRC, SUB, CA1, and CA2/3/DG) and group (old/young) as fixed effects, random intercepts for each participant and random slopes for ROI for each participant to test this possibility. There were no significant differences across the four types of trials in CA1 in younger adults (the Young-Adult atlas: *ps* > 0.903; the Old-Adult atlas: *ps* > 0.579). This control analysis ruled out that the possibility that the remapping effect that found in younger adults were driven by higher within-city perceptual information.

<u>*More variable neural representation for the same environment in older adults in CA1 (results from the* Old-Adult atlas*):*</u> Again, we re-examined standard deviation of neural representation using the Old-Adult atlas to ensure the robustness of our findings. The MPS of older adults showed a wider distribution of values than younger adults, indicating a broader spread of activation patterns within a city (Fig. S2b). Direct comparison of younger versus older adult within-city MPS revealed a larger standard deviation for within-city MPS in CA1 of older individuals ($t(45) = 2.993$, $p = 0.004$, Cohen's D = 0.875; Fig. S3b). The Bayes factor calculated for the standard deviation for within-city MPS indicated strong evidence against the null hypothesis ($BF_{10} = 9.201$, with strong evidence). This effect could not be accounted for by differences in performance when using spatial distance discrimination performance as covariate in a linear regression model ($R^2_{adjusted} = 0.129$, $F(2, 44) = 4.399$, $p = 0.018$).

The relationship between variable neural representation for the same environment and spatial distance discrimination performance (results from both Young-Adult and Old-Adult atlas): We also found that the higher standard deviation for within-city MPS in CA1, the worse spatial distance discrimination performance (Pearson correlation: Young-Adult atlas: $r(45) = -0.416$, $p = 0.004$, Spearman correlation: $r(45) = -0.429$, $p = 0.003$, $BF_{10} = 10.975$, with strong evidence; Old-Adult atlas: Pearson correlation: $r(45) = -0.225$, $p = 0.129$; Spearman correlation: $r(45) = -0.335$, $p = 0.022$, Fig. S4). To further account for potential outliers and enhance the reliability of our analysis, we performed robust linear regression using the "robustfit" function in MATLAB, and the results still persisted (Young-Adult atlas: $\beta = -11.299$, $t(45) = -3.732$, $p < 0.001$; Old-Adult atlas: $\beta = -4.986$, $t(45) = -1.920$, $p = 0.061$). This correlation did not survive when adding age as covariate (Young-Adult atlas: $r_{partial}(45) = -0.091$, $p = 0.559$; Old-Adult atlas: Pearson correlation: $r_{partial}(45) = -0.005$, $p = 0.973$; Spearman correlation: $r_{partial}(45) = -0.188$, $p = 0.210$). This finding is consistent with an age-variant effect such that older adults showed a higher variance in within-city MPS, regardless of performance.

<u>*Control analysis after matching number of trials for calculating standard deviation of within-city MPS in CA1 (results from both atlases):*</u> As the sample size increases, the standard deviation decreases. In the current study, younger adults had more trial pairs than older adults, which could lead to a decreased SD. To exclude the possibility that age-differences in SD were driven by unequal number of trial pairs, we conducted a control analysis to match the trial pairs by randomly resampling trial pairs of each participant with the smallest number of trial pairs for the within-city condition. We repeated this procedure 5000 times. When applying the analysis on the Young-Adult atlas, the results showed that the standard deviation of within-city MPS in CA1 was higher in older adults than in younger adults ($t(45) = 6.245$ , $p < 0.001$, FDR corrected; Cohen's *D* = 1.826; $BF_{01} = 79431.998$, with extremely strong evidence ). When applying the same analysis on the Old-Adult atlas, the results again showed that standard deviation of within-city

MPS in CA1 was higher in older adults than younger adults (t(45) = 5.534, p < 0.001, FDR corrected; Cohen's *D* = 1.618, $BF_{01}$ = 8792.887, with extremely strong evidence).

<u>*More variable neural representation for the same environment in other MTL ROIs (results from both atlases).*</u> Besides CA1, we also observed lower standard deviation for within-city MPS in other MTL ROIs when performing the analysis using both atlases, including CA2/3/DG (the Young-Adult atlas: t(45) = 3.828, p = 0.004, Cohen's *D* = 1.119, $BF_{10}$ = 67.518, with very strong evidence; the Old-Adult atlas: t(45) = 3.603, p = 0.003, Cohen's *D* = 1.053, $BF_{10}$ = 38.25, with very strong evidence; FDR corrected), ERC (the Young-Adult atlas: t(45) = 3.4, p = 0.003, Cohen's *D* = 0.994; $BF_{10}$ = 23.312, with strong evidence; the Old-Adult atlas: t(45) = 2.338, p = 0.024, Cohen's *D* = 0.684; $BF_{10}$ = 2.508, with anecdotal evidence; FDR corrected), PRC (the Young-Adult atlas: t(45) = 2.24, p = 0.036, Cohen's *D* = 0.656; $BF_{10}$ = 2.216, with anecdotal evidence; the Old-Adult atlas: t(45) = 4.443, p = 0.003, Cohen's *D* = 1.299; $BF_{10}$ = 354.762, with extreme evidence; FDR corrected) for older adults than younger adults. Besides, the SD of within-city PS in PRC was negatively correlated with individuals' spatial distance discrimination performance (the Young-Adult atlas: Pearson correlation: r(45) = -0.449, p < 0.01; Spearman correlation r(45) = -0.552, p < 0.001, $BF_{10}$ = 23.585, with strong evidence; the Old-Adult atlas: Pearson correlation: r(45) = -0.534, p < 0.001;Spearman correlation: r(45) = -0.684, p < 0.001, $BF_{10}$ = 254.697, with extremely strong evidence, Fig. S4). To further account for potential outliers and enhance the reliability of our analysis, we performed robust linear regression using the "robustfit" function in MATLAB, and the results still persisted (Young-Adult atlas: β = -5.186, t(45) = -4.064, p < 0.001; Old-Adult atlas: β = -12.685, t(45) = -9.497, p < 0.001). But this correlation between the SD of within-city PS in PRC and individuals' spatial distance discrimination performance was not significant after controlling for age, gender and PRC volume (the Young-Adult atlas: $r_{partial}$(45) = -0.159, p = 0.303; the Old-Adult atlas: $r_{partial}$ (45) = -0.285, p = 0.061).

**Supplementary Note 3: Neural remapping and spatial distance discrimination are age-invariant**

<u>*The age-invariant relationship between neural remapping index and spatial distance discrimination performance (results from the Old-Adult atlas)*</u>: We found that the neural remapping index in CA1 was positively correlated with participant spatial distance discrimination performance as measured by the percent of successfully retrieved distances of stores (r(45) = 0.444, p = 0.002, $BF_{10}$ = 21.000, with strong evidence, Fig. S5b). This correlation was age-invariant (after controlling for age, gender, and CA1 volume, the correlation persisted, i.e., $r_{partial}$(45) = 0.418, p = 0.005). We did not find any correlation between the neural remapping index in other MTL subregions (ps > 0.126, FDR corrected, Fig. S5b). Together, these findings suggest that better performing participants – regardless of age – showed greater differences in the CA1 remapping index.

**Supplementary Note 4: Differences in the neural remapping index related to spatial distance discrimination are mediated by alterations in input to CA1**

<u>*Increased interregional informational connectivity in older adults (results from the Old-Adult atlas):*</u> We obtained similar results when we assayed interregional informational connectivity using the Old-Adult atlas. We found that older adults showed increased interregional informational connectivity between ERC-PRC (t(45) = 1.905, p = 0.002, FDR corrected; Cohen's D = 1.214, $BF_{10}$ = 159.888, with extremely strong evidence), ERC-CA1 (t(45) = 3.475, p = 0.007, FDR corrected; Cohen's *D* = 1.016, $BF_{10}$ = 27.885, with strong evidence), ERC-CA2/3/DG (t(45) = 2.732, p = 0.022, FDR corrected; Cohen's *D* = 0.799, $BF_{10}$ = 5.309, with moderate evidence), PRC-SUB (t(45) = 2.758, p = 0.022, FDR corrected; Cohen's *D* = 0.806, $BF_{10}$ = 5.599, with moderate evidence), PRC-CA1 (t(45) = 3.037, p = 0.016, FDR corrected;

Cohen's $D$ = 0.888, $BF_{10}$ = 10.118, with strong evidence), SUB-CA1 ($t(45)$ = 2.499, p = 0.032, FDR corrected; Cohen's $D$ = 0.730, $BF_{10}$ = 3.366, with moderate evidence) and CA2/3/DG-CA1 ($t(45)$ = 2.24, p = 0.05, FDR corrected; Cohen's $D$ = 0.654, $BF_{10}$ = 2.103, with anecdotal evidence Fig. S6c). These findings suggested that the interactions between different subfields were more similar, on average, in older than younger adults. Here, we specifically focused on the connections between input areas (i.e., PHC, PRC, ERC, SUB and CA2/3/DG) to CA1 (Fig. 3c). We calculated the interregional information connectivity by averaging across four selected connections: ERC-CA1, PRC-CA1, SUB-CA1 and CA2/3/DG-CA1 (see Methods). We found a negative correlation between the interregional informational connectivity and spatial distance discrimination performance ($r(45)$ = -0.450, p = 0.002, $BF_{10}$ = 23.925, with strong evidence, Fig. S6f) as well as interregional informational connectivity and the neural remapping index ($r(45)$ = -0.246, p = 0.096, $BF_{10}$ = 0.700, Fig. S6g). This suggests that a higher correlation between patterns in different subfields related to worse spatial distance discrimination and a lower neural remapping index.

*Differences in the neural remapping index related to spatial distance discrimination are mediated by alterations in input to CA1 (results from the Old-Adult atlas):* To attempt to better understand the relationship between interregional informational connectivity patterns, the neural remapping index, and spatial distance discrimination, we performed a mediation analysis. The model indicated that interregional informational connectivity (Table S6) mediated the relationship between the neural remapping index in CA1 and spatial distance discrimination after controlling for CA1 volume and gender (Fig. S4d). Notably, when we entered age as a covariate, however, the mediation effect was no longer significant, suggesting that the interregional informational connectivity mediation effect was age-variant (Table S6). We also examined the informational connectivity between PRC-ERC, which has an indirect connection with the CA1, and the results suggest that the PRC-ERC informational connectivity also contributed to relationship between the neural remapping index effect in CA1 and spatial distance discrimination after controlling for CA1 volume and gender. This effect did not survive after adding age as a covariate (Table S10). Together, these findings suggest that differences in interregional informational connectivity contribute to age-variant differences in the neural remapping index.

*Differences in the stability of patterns for the same environment related to spatial distance discrimination are mediated by alterations in input to CA1 (results from the Young-Adult atlas):* Here, we quantified interregional informational connectivity using the values of the correlations of within-city MPS between one subfield and another. We found that older adults showed increased interregional informational connectivity between PRC-ERC ($t(45)$ = 3.756, p = 0.003, FDR corrected; Cohen's D = 1.098, $BF_{10}$ = 56.208, with very strong evidence), ERC-SUB ($t(45)$ = 2.428, p = 0.039, FDR corrected; Cohen's $D$ = 0.710, $BF_{10}$ = 2.950, with anecdotal evidence), ERC-CA1 ($t(45)$ = 3.159, p = 0.011, FDR corrected; Cohen's $D$ = 0.923, $BF_{10}$ = 13.277, with strong evidence), ERC-CA23DG ($t(45)$ = 2.876, p = 0.018, FDR corrected; Cohen's $D$ = 0.841, $BF_{10}$ = 7.158, with moderate evidence), PRC-SUB ($t(45)$ = 2.761, p = 0.020, FDR corrected; Cohen's $D$ = 0.807, $BF_{10}$ = 5.636, with moderate evidence), SUB-CA1 ($t(45)$ = 3.913, p = 0.003, FDR corrected; Cohen's $D$ = 1.144, $BF_{10}$ = 84.264, with very strong evidence). Here, we specifically focused on the connections between input areas (i.e., PHC, PRC, ERC, SUB and CA23DG) to CA1. We calculated the interregional information connectivity by averaging across three selected connections: PHC-CA1, ERC-CA1, and SUB-CA1, which were selected based on the p-value of the contrast of group difference (see Methods). To attempt to better understand the relationship between interregional informational connectivity, the stability of patterns for the same environment, and spatial distance discrimination, we performed a mediation analysis. The model indicated that informational connectivity (Table S11) mediated the relationship between the standard deviation of within-city MPS in CA1 and spatial distance discrimination performance after controlling for CA1 volume and gender. Notably, when we entered age as a covariate, however, the mediation effect was no longer significant, suggesting that the interregional heterogeneity mediation effect was age-variant (Table S11). Similar results were found when using other connectivity patterns to CA1 (ERC-CA1, SUB-CA1 and CA2/3/DG-CA1); See Table S12. Together, these findings suggest that differences in the input to CA1 contribute to age-variant differences in the stability of patterns for the same environment.

*Differences in the stability of patterns for the same environment related to spatial distance discrimination are mediated by alterations in input to CA1 (results from the Old-Adult atlas):* Here, we quantified interregional informational connectivity using the values of the correlations of within-city MPS between one subfield and another. We found that older adults showed increased interregional informational connectivity between ERC-PRC ($t(45) = 5.032$, $p < 0.001$, FDR corrected; Cohen's $D = 1.471$, $BF_{10} = 1942.889$, with extremely strong evidence), ERC-SUB ($t(45) = 2.436$, $p = 0.038$, FDR corrected; Cohen's $D = 0.712$, $BF_{10} = 2.994$, with anecdotal evidence), ERC-CA1 ($t(45) = 3.063$, $p = 0.015$, FDR corrected; Cohen's $D = 0.895$, $BF_{10} = 10.716$, with strong evidence), ERC-CA23DG ($t(45) = 2.663$, $p = 0.032$, FDR corrected; Cohen's $D = 0.778$, $BF_{10} = 4.620$, with moderate evidence), PRC-SUB ($t(45) = 2.491$, $p = 0.038$, FDR corrected; Cohen's $D = 0.728$, $BF_{10} = 3.318$, with moderate evidence), PRC-CA1 ($t(45) = 3.144$, $p = 0.015$, FDR corrected; Cohen's $D = 0.919$, $BF_{10} = 12.864$, with strong evidence). Here, we specifically focused on the connections between input areas (i.e., PHC, PRC, ERC, SUB and CA23DG) to CA1. We calculated the interregional information connectivity by averaging across three selected connections: ERC-CA1, PRC-CA1, and SUB-CA1, which were selected based on the p-value ($p < 0.05$, uncorrected) of the contrast of group difference (see Methods). To attempt to better understand the relationship between interregional informational connectivity, the stability of patterns for the same environment, and spatial distance discrimination, we performed a mediation analysis. The model indicated that informational connectivity (Table S13) mediated the relationship between the standard deviation of within-city MPS in CA1 and spatial distance discrimination performance after controlling for CA1 volume and gender. Notably, when we entered age as a covariate, however, the mediation effect was no longer significant, suggesting that the interregional informational connectivity mediation effect was age-variant (Table S13). Similar results were found when using other connectivity patterns to CA1 (ERC-CA1, SUB-CA1 and CA2/3/DG-CA1); See Table S14. Together, these findings suggest that differences in the input to CA1 contribute to age-variant differences in the stability of patterns for the same environment.

### Supplementary Note 5: Correlations between the neural remapping index and spatial distance discrimination are mediated by reduced CA1 neural consistency

Age-invariant differentiation relates to spatial distance discrimination performance and is mediated by reduced dimensionality of CA1 *(results from the Old-Adult atlas)*. We obtained similar results when we applied the principal components analysis using the Old-Adult atlas. The results showed that the dimensionality of CA1 was significantly lower in older adults compared to younger adults ($t(45) = 4.288$, $p < 0.001$, Cohen's D = 1.253, $BF_{10} = 230.747$,with extremely strong evidence; Fig. S7d) after matching the number of voxels by selecting the least number of voxels in the whole group. Furthermore, we found that the dimensionality of CA1 was positively associated with neural remapping index ($r(45) = 0.535$, $p < 0.001$, $BF_{10} = 258.841$, with extremely strong evidence; Fig. S7c) and spatial distance discrimination ($r(45) = 0.805$, $p < 0.001$, $BF_{10} = 1,081,000,000$, with extremely strong evidence; Fig. S7b). These effects were significant even after controlling for age, gender, and CA1 volume, even after controlling for age, gender and CA1 volume (neural remapping index: $r_{partial}(45) = 0.496$, $p < 0.001$; spatial distance discrimination: $r_{partial}(45) = 0.737$, $p < 0.001$). As discussed earlier, we also found a correlation between spatial distance discrimination and the neural remapping index in CA1 (Fig. S5b and Supplementary Note 3). Thus, we then ran an analysis to determine if the number of extracted principal components in CA1 mediates this effect. This analysis indicated that the dimensionality of CA1 did mediate the correlation between the neural remapping index and spatial distance discrimination, even after controlling for CA1 volume and gender (Table S16). The mediation effect remained significant when age was included as a covariate, suggesting that these effects were age-invariant (Table S16). These findings suggest that reduced fidelity of signals within CA1 mediated the relationship between the neural remapping index and participants' memories for the distances of landmarks to each other in each city. Therefore, increased neural differentiation between different cities was related to better performance, based on the fidelity of signals within CA1 – regardless of age.

*Control for voxel number when estimating the dimensionality (results from both atlases).* Since the number of principal components depends on the number of voxels involved in PCA, we performed two different control analyses to rule out the possibility that the age-related differences were driven by voxel number. First, we used top 50 voxels based on each participant's tSNR level to examine the dimensionality. The results revealed that the dimensionality of CA1 was significantly lower in older adults compared to younger adults (Young-Adult atlas: $t(45) = 4.075$, $p < 0.001$, Cohen's $D = 1.191$, $BF_{10} = 650,000,000$, with extremely strong evidence; Old-Adult atlas: $t(45) = 3.839$, $p < 0.001$; Cohen's $D = 1.122$, $BF_{10} = 628,000,000$, with extremely strong evidence;). Second, we performed the PCA using all voxels within each individual CA1 and again found lower dimensionality in older adults than in younger adults (the Young-Adult atlas: $t(45) = 4.867$, $p < 0.001$, Cohen's $D = 1.422$, $BF_{10} = 311900$, with extremely strong evidence;; the Old-Adult atlas: $t(45) = 4.486$, $p < 0.001$, Cohen's $D = 1.311$, $BF_{10} = 654,600,000$, with extremely strong evidence;).

*Control for trial numbers when estimating the dimensionality (results from both atlases).* The dimensionality estimation was based on the N-voxel by N-voxel functional connectivity matrix derived from the Pearson correlation of t-statistics series of correct trials. One possibility is that this functional connectivity matrix of older adults was biased by fewer corrected trials than younger adults. To investigate this, we conducted a control analysis by excluding 5 participants with the least number of trials from the old group and 7 participants with the most number of trials from the young group to make the number of correct trials between the two group comparable ($Mean_{OA} = 85$, $Mean_{YA} = 95$, $t(33) = 1.92$, $p = 0.064$; Cohen's $D = 0.649$). Then we recompared the dimensionality between the two sub-groups ($N_{OA} = 17$, $N_{YA}=18$) while matching the number of voxels for the whole group. The results showed that the dimensionality of CA1 was lower in older adults than younger adults (the Young-Adult atlas: $t(33) = 2.077$, $p = 0.046$; Cohen's $D = 0.702$, $BF_{10} = 1.651$, with anecdotal evidence ,the Old-Adult atlas: $t(33) = 2.024$, $p = 0.051$; Cohen's $D = 0.685$, $BF_{10} = 1.526$, with anecdotal evidence). This result did not change when we used top 50 voxels based on tSNR (the Young-Adult atlas: $t(33) = 2.280$, $p = 0.029$; Cohen's $D = 0.771$, $BF_{10} = 2.268$, with anecdotal evidence; the Old-Adult atlas: $t(33) = 1.94$, $p = 0.06$; Cohen's $D = 0.656$, $BF_{10} = 1.350$, with anecdotal evidence) or using all voxels within CA1 (the Young-Adult atlas: $t(33) = 2.733$, $p = 0.01$, Cohen's $D = 0.917$, $BF_{10} = 4.481$, with moderate evidence; the Old-Adult atlas: $t(33) = 2.16$ , $p = 0.038$, Cohen's $D = 0.731$, $BF_{10} = 1.875$, with anecdotal evidence).

*Reduced dimensionality in other hippocampal subfields (results from both atlas).* We also estimated the dimensionality of neural signals by performing PCA to determine whether the fidelity of signals within hippocampus subfields also contributed to variance in spatial distance discrimination performance. The results showed that the dimensionality of PHC, PRC, ERC, SUB and CA23DG was significantly lower in older adults compared to younger adults (Young-Adult atlas: $ts(45) > 4.130$, Cohen's $Ds > 1.207$, $ps < 0.001$, $BF_{10} > 149.933$, with extreme strong evidence; Old-Adult atlas: ts(45) > 3.994, Cohen's *Ds* > 1.168, $p < 0.001$, $BF_{10}$ > 104.250, with extreme strong evidence).

We found that dimensionality of PHC, PRC, ERC, SUB and CA23DG was positively associated with spatial distance discrimination (Young-Adult atlas: rs(45) > 0.798, ps < 0.001, $BF_{10}$ > 574,100,000, with extreme evidence; Old-Adult atlas: rs(45) > 0.759, ps < 0.001, $BF_{10}$ > 20,060,000, with extreme strong evidence). However, none of these subregions' dimensionality was correlated with their neural remapping index (Young-Adult atlas: rs(45) < 0.238, ps > 0.08, $BF_{01}$ > 1.566; Old-Adult atlas: ps > 0.106, $BF_{01}$ > 0.667). We also did not find any evidence of mediation effects in other MTL subregions.

**Supplementary Note 6: Functional connectivity between MTL ROIs.** We assessed the functional connectivity between MTL sub-regions by using a fMRI t-statistics series approach. This is analogous to the beta-time series method but with the beta-values transformed to t-statistics(1). Briefly, we generated a time-series of single-trial t-statistics from GLMs (see Single-trial response estimates) and averaged these across all voxels within each ROI. Note, we only used correct trials and also removed trials that included any volume exceeding the framewise displacement threshold of 0.5 mm from the t-statistics series. We then computed the Pearson correlation coefficient between the t-statistics series of all ROI pairs, resulting in a correlation matrix that describes the strength of the functional relationship between any two regions for each participant. Our analysis did not reveal any significant differences between the two groups in terms of the functional connectivity of the MTL ROIs (the Young-Adult atlas: *ps* > 0.636; the Old-Adult atlas: *ps* > 0.294; FDR corrected).

### Supporting Figures

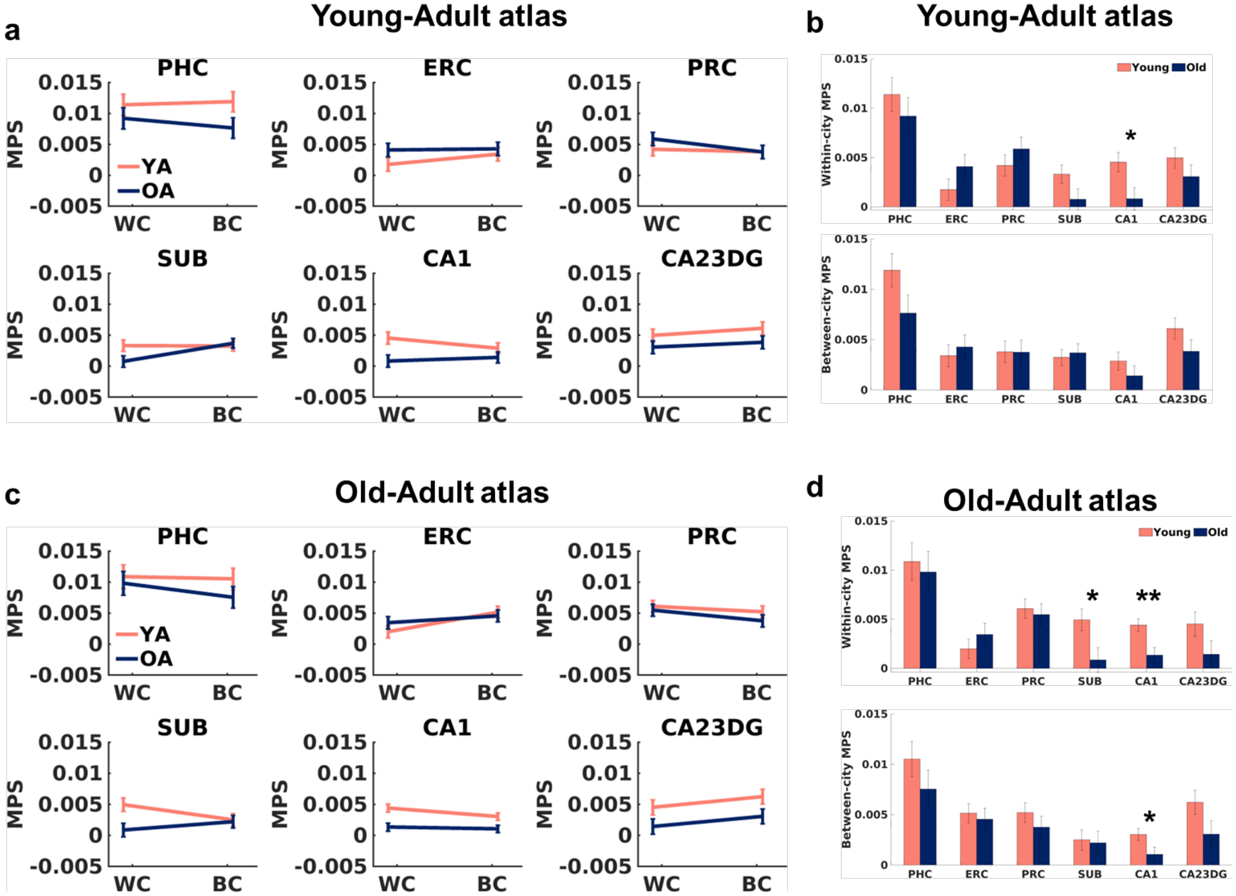


**Fig. S1: Results from a linear mixed effects model with *condition* (within-city/between-city), *ROI* (PHC, ERC, PRC, SUB, CA1, and CA2/3/DG, segmentation based on the Young-Adult and the Old-Adult atlas) and *group* (old/young) as fixed effects, random intercepts for each participant and random slopes for *condition* and random slopes for *ROI* for each participant using both Young-Adult and Old-Adult atlases.** The data shown here are estimated marginal means from mixed effects models. WC: within-city, BC: between-city, MPS: multivariate pattern similarity. Each individual dot represents data from an individual participant. All data reflect n = 25 for YA, and n = 22 for OA independent participants. YA: younger adults (red), OA: older adults (dark blue). *p < 0.05, **p < 0.01. p-value was FDR corrected.

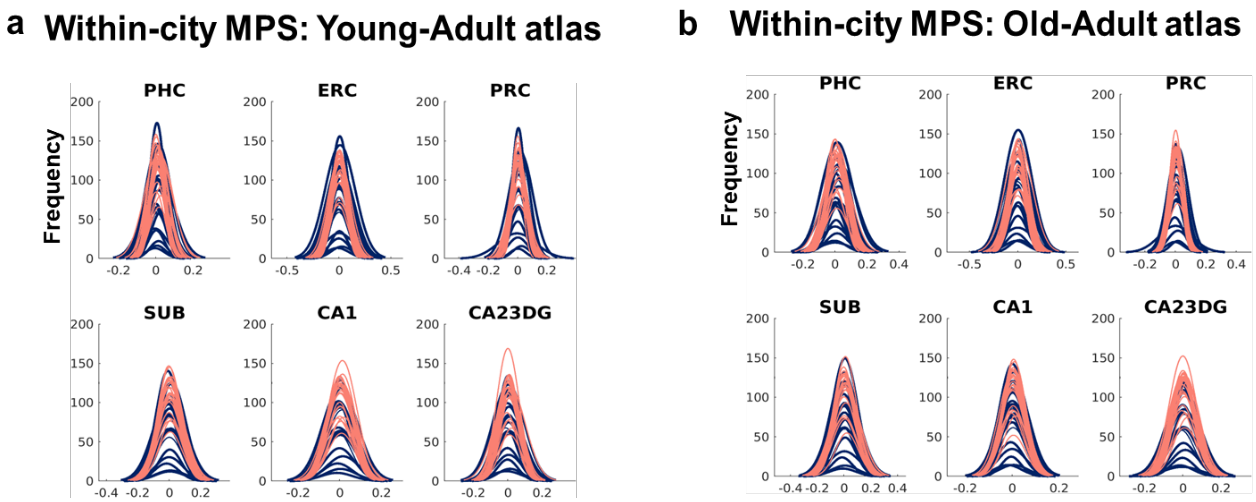


**Fig. S2: The within-city MPS of older adults showed a wider distribution of values than younger adults.** a. Distribution of within-city MPS of each ROI (results from the Young-Adult atlas). b. Distribution of within-city MPS of each ROI (results from the Old-Adult atlas). Each individual line represents data from an individual participant. All data reflect n = 25 for YA, and n = 22 for OA independent participants. MPS: multivariate pattern similarity. YA: younger adults (red), OA: older adults (dark blue).

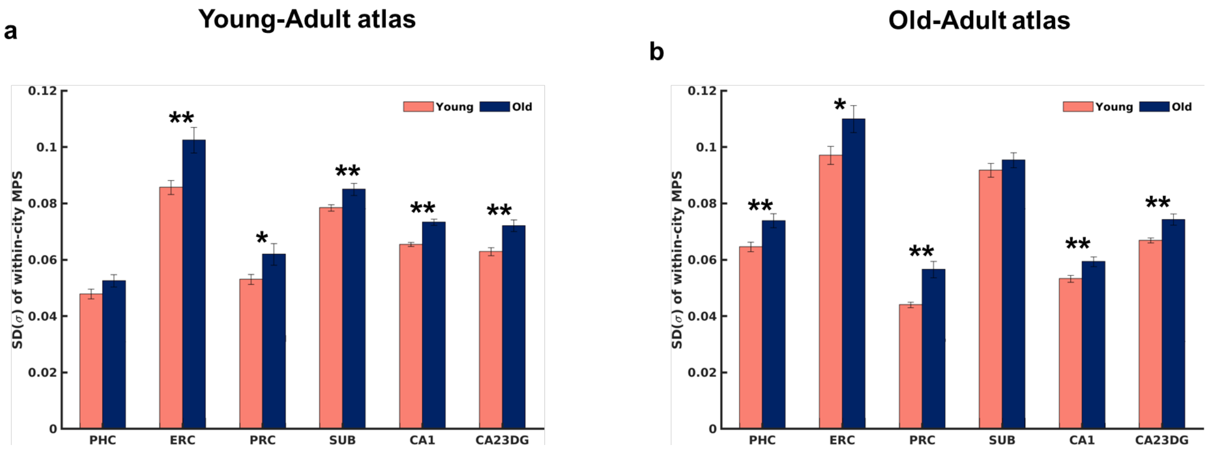


**Fig. S3: Higher standard deviation of within-city MPS was found for older than younger adults on both atlases.** a. Standard deviation of within-city MPS of each ROI (results from the Young-Adult atlas). b. Standard deviation of within-city MPS of each ROI (results from the Old-Adult atlas). MPS: multivariate pattern similarity. Each individual dot represents data from an individual participant. All data reflect n = 25 for YA, and n = 22 for OA independent participants. YA: younger adults (red), OA: older adults (dark blue). *p < 0.05, **p < 0.01. p-value was FDR corrected.

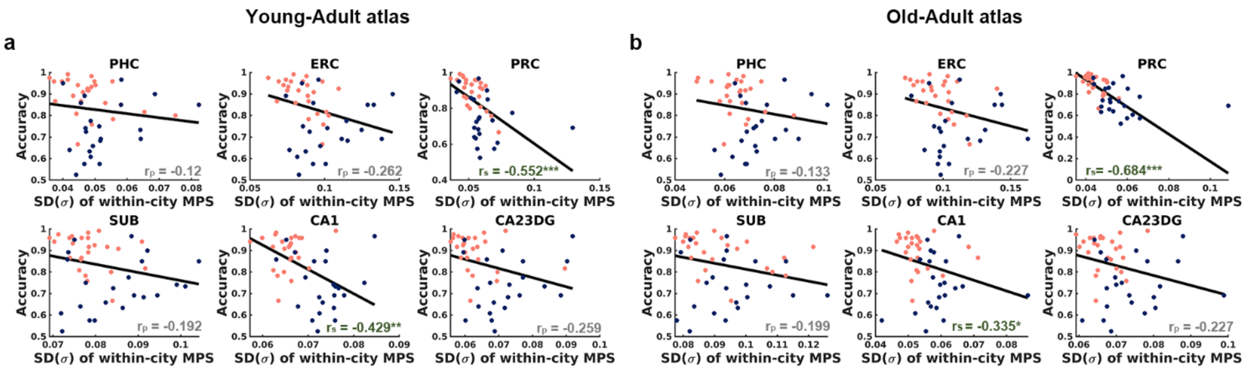


**Fig. S4: The correlation between spatial distance discrimination and standard deviation of within-city MPS in each ROI for each atlas.** Each individual dot represents data from an individual participant. All the trend lines were fitted using robust linear regression. All data reflect n = 25 for YA, and n = 22 for OA independent participants**.** $r_p$: Pearson correlation coefficient; $r_s$: Spearman correlation coefficient. YA: younger adults (red), OA: older adults (dark blue). *p < 0.05, **p < 0.01, ***p < 0.001.

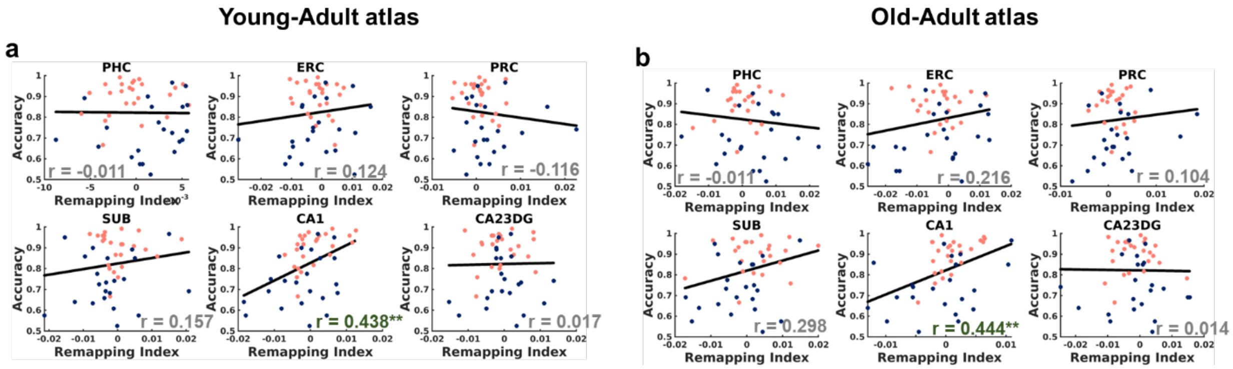


**Fig. S5: The correlation between remapping index and accuracy in each ROI for each atlas**. Each individual dot represents data from an individual participant. All the trend lines were fitted using robust linear regression. All data reflect n = 25 for YA, and n = 22 for OA independent participants. YA: younger adults (red), OA: older adults (dark blue). **p < 0.01.

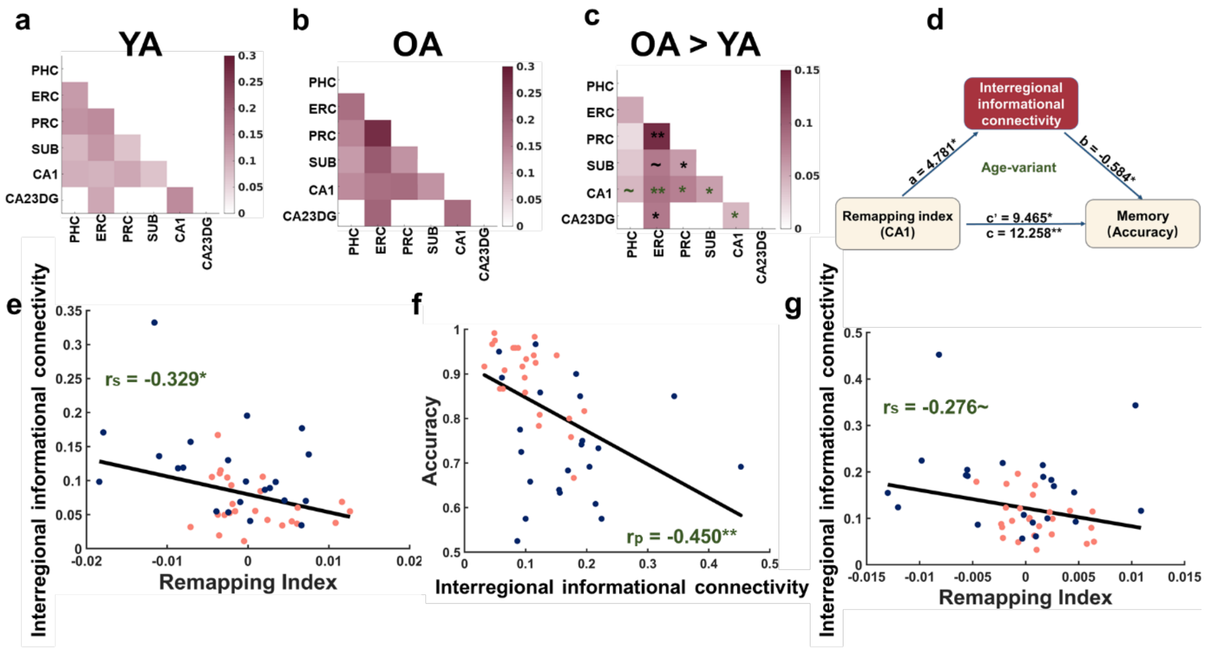


**Fig. S6: Differences in the remapping index related to spatial distance discrimination performance are mediated by alterations in input to CA1.** a. Interregional informational connectivity in younger adults (results from the Old-Adult atlas). b. Interregional informational connectivity in older adults (results from the Old-Adult atlas). c. Reduced interregional informational connectivity in older adults from subfields with direct input to CA1 (green asterisk, results from the Old-Adult atlas). d. Interregional informational connectivity mediated the relationship between the neural remapping index in CA1 and spatial distance discrimination performance after controlling for CA1 volume and gender, but this mediation effect was no longer significant after adding age as covariate (results from the Old-Adult atlas). e. The interregional informational connectivity was negatively associated with neural remapping index (i.e., within minus between city MPS) in CA1 (results from the Young-Adult atlas). f. The interregional informational connectivity was negatively associated with spatial distance discrimination performance (results from the Old-Adult atlas). g. The interregional informational connectivity was negatively associated with neural remapping index (i.e., within minus between city MPS) in CA1 (result from the Old-Adult atlas). All the trend lines were fitted using robust linear regression. Each individual dot represents data from an individual participant. All data reflect n = 25 for YA, and n = 22 for OA independent participants. ~$p < 0.1$, *$p < 0.05$, **$p < 0.01$, ***$p < 0.001$. $r_p$: Pearson correlation coefficient; $r_s$: Spearman correlation coefficient. YA: younger adults (red), OA: older adults (dark blue). p-value was FDR corrected.

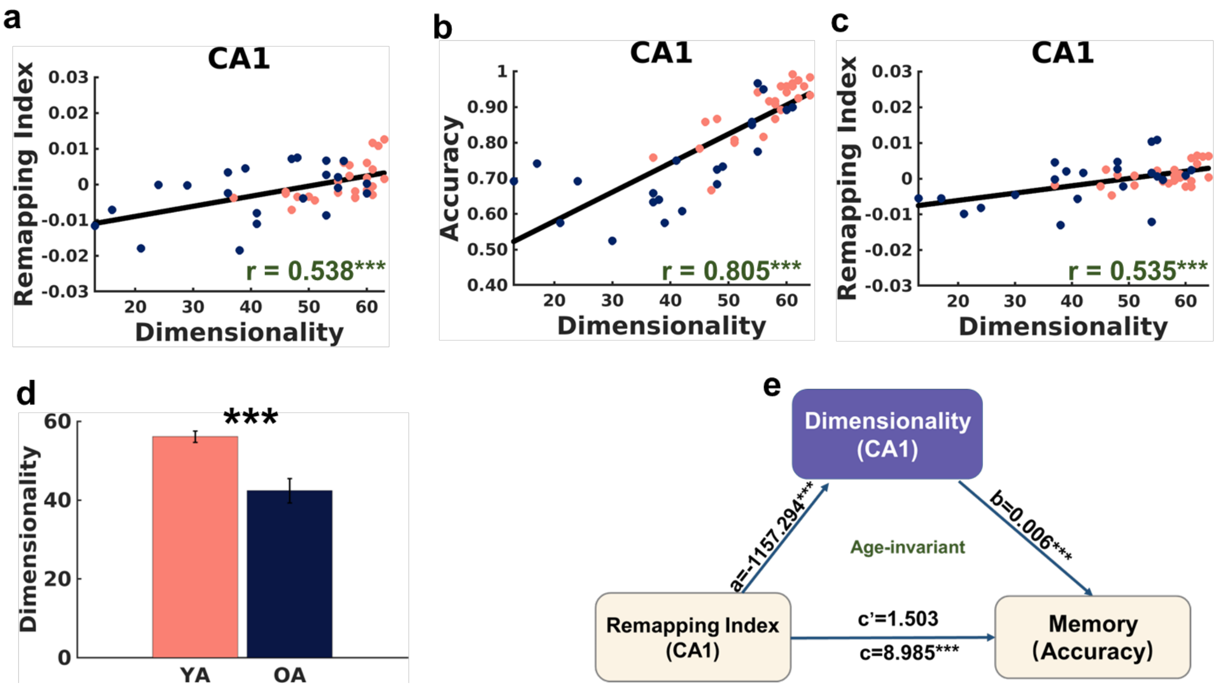


**Fig. S7: Correlations between the remapping index and spatial distance discrimination performance are mediated by reduced CA1 neural consistency.** a. The consistency ("dimensionality") was positively associated with neural remapping index (within-city MPS – between-city MPS; result from the Young-Adult atlas). b. The dimensionality of CA1 was positively associated with spatial distance discrimination performance (result from the Old-Adult atlas). c. The dimensionality of CA1 was positively associated with neural remapping index (within-city MPS – between-city MPS; result from the Old-Adult atlas). d. Reduced CA1 neural consistency in older adults (result from the Old-Adult atlas). e. The dimensionality of CA1 fully mediated the effect of the neural remapping index on spatial distance discrimination performance, even after controlling for age, gender and CA1 volume (results from the Old-Adult atlas). Error bars represent the standardized errors of the means. Each individual dot represents data from an individual participant. All the trend lines were fitted using robust linear regression. All data reflect n = 25 for YA, and n = 22 for OA independent participants. $^{**}p < 0.01$, $^{***}p < 0.001$. YA: younger adults (red), OA: older adults (dark blue).

### Supporting Tables

Table S1: Results from linear mixed effects models predicting pattern. Models include random intercepts for each participant, random slopes for *Condition* (within-city/between-city), and random slopes for *ROI* for each participant (i.e., PHC, ERC, PRC, SUB, CA1 and CA2/3/DG; segmentation based on the Young-Adult atlas).

| Fixed Effects | NumDF | DenDF | F-value | *p-value* |
|---|---|---|---|---|
| Condition | 1 | 42 | 0.136 | 0.714 |
| ROI | 5 | 45 | 10.135 | 0.000*** |
| Group | 1 | 44 | 1.011 | 0.320 |
| Condition*ROI | 5 | 974483 | 8.081 | 0.000*** |
| Condition*Group | 1 | 42 | 0.006 | 0.941 |
| ROI*Group | 5 | 45 | 1.679 | 0.159 |
| Condition*ROI*Group | 5 | 974483 | 8.232 | 0.000*** |

All data reflect n = 25 for YA, and n = 22 for OA independent participants. NumDF: degrees of freedom for the numerator, DenDF: degrees of freedom for the denominator. ***p < 0.001.

Table S2: Results from linear mixed effects models predicting pattern similarity after controlling for univariate activation levels, activation variance, spatial distance discrimination performance, reaction time, and gender. Models include random intercepts for each participant, random slopes for *Condition* (within-city/between-city), and random slopes for *ROI* for each participant (i.e., PHC, ERC, PRC, SUB, CA1, and CA2/3/DG; segmentation based on the Young-Adult atlas).

| Fixed effects | anova | DenDF | F-value | *p-value* |
|---|---|---|---|---|
| Condition | 1 | 42 | 0.173 | 0.679 |
| ROI | 5 | 55 | 4.678 | 0.001** |
| Group | 1 | 45 | 3.826 | 0.057 |
| $ACT_{mean}$ | 1 | 850594 | 194.110 | 0.000*** |
| $ACT_{sd}$ | 1 | 684728 | 20.213 | 0.000*** |
| ROI Volume | 1 | 56 | 4.729 | 0.034* |
| Performance | 1 | 41 | 0.839 | 0.365 |
| RT | 1 | 40 | 4.169 | 0.048* |
| Gender | 1 | 40 | 4.952 | 0.032* |
| Condition*ROI | 5 | 974483 | 8.070 | 0.000*** |
| Condition*Group | 1 | 42 | 0.004 | 0.950 |
| ROI*Group | 5 | 46 | 1.516 | 0.203 |
| Condition*ROI*Group | 5 | 974483 | 8.231 | 0.000*** |

All data reflect n = 25 for YA, and n = 22 for OA independent participants. NumDF: degrees of freedom for the numerator, DenDF: degrees of freedom for the denominator. ACTmean: mean activation levels, ACTSD: activation variance, RT: reaction time. *p < 0.05, **p< 0.01, ***p < 0.001.

Table S3: Results from linear mixed effects models predicting pattern similarity. Models include random intercepts for each participant, random slopes for Condition (within-city/between-city), and random slopes for *ROI* for each participant (i.e., PHC, ERC, PRC, SUB, CA1 and CA2/3/DG; segmentation based on the Old-Adult atlas).

| Fixed Effects | NumDF | DenDF | F-value | *p-value* |
| --- | --- | --- | --- | --- |
| Condition | 1 | 44 | 0.004 | 0.949 |
| ROI | 5 | 45 | 8.636 | 0.000*** |
| Group | 1 | 42 | 2.617 | 0.113 |
| Condition*ROI | 5 | 974474 | 14.417 | 0.000*** |
| Condition*Group | 1 | 44 | 0.000 | 0.994 |
| ROI*Group | 5 | 45 | 1.430 | 0.232 |
| Condition*ROI*Group | 5 | 974474 | 7.412 | 0.000*** |

All data reflect n = 25 for YA, and n = 22 for OA independent participants. NumDF: degrees of freedom for the numerator, DenDF: degrees of freedom for the denominator. ***p < 0.001.

Table S4: Results from linear mixed effects models predicting pattern similarity after controlling for univariate activation levels, activation variance, spatial distance discrimination performance, reaction time and gender. Models include random intercepts for each participant, and random slopes for *ROI* for each participant (i.e., PHC, ERC, PRC, SUB, CA1 and CA2/3/DG; segmentation based on the Old-Adult atlas).

| Effects | NumDF | DenDF | F-value | *p-value* |
|---|---|---|---|---|
| Condition | 1 | 974552 | 6.2763 | 0.012* |
| ROI | 5 | 56 | 3.758 | 0.005** |
| Group | 1 | 51 | 9.798 | 0.003** |
| $ACT_{mean}$ | 1 | 641247 | 230.506 | 0.000*** |
| $ACT_{sd}$ | 1 | 349488 | 22.836 | 0.000*** |
| Voxel number | 1 | 56 | 0.154 | 0.696 |
| Performance | 1 | 47 | 3.981 | 0.052 |
| RT | 1 | 40 | 3.733 | 0.06 |
| Gender | 1 | 37 | 6.614 | 0.014* |
| Condition*ROI | 5 | 974518 | 14.983 | 0.000*** |
| Condition*Group | 1 | 974552 | 6.001 | 0.014* |
| ROI*Group | 5 | 46 | 1.371 | 0.252 |
| Condition*ROI*Group | 5 | 974518 | 7.829 | 0.000*** |

All data reflect n = 25 for YA, and n = 22 for OA independent participants. NumDF: degrees of freedom for the numerator, DenDF: degrees of freedom for the denominator. ACTmean: mean activation levels, ACTSD: activation variance, RT: reaction time. *p < 0.05, **p< 0.01, ***p < 0.001.

Table S5: Mediation analysis using interregional informational connectivity as a mediator, calculated by averaging interregional informational connectivity across unique edges from the Young-Adult atlas (including PHC-CA1, ERC-CA1, PRC-CA1, SUB-CA1 and CA2/3D/G-CA1), with the neural remapping index of CA1 as an independent variable and spatial distance discrimination performance as a dependent variable (results from the Young-Adult atlas).

| Covariate | c<br>Total effect | a | b | a*b<br>Mediation effect | a*b<br>(95% BootCI) | c'<br>Direct effect | Conclusion |
|---|---|---|---|---|---|---|---|
| CA1 Volume and Gender | 9.314*** | -3.637** | -0.750* | 2.728 | 0.840~5.406 | 6.586* | Partial mediation |
| CA1 Volume, Gender and Age | 6.592** | -2.897* | -0.449 | 1.299 | -0.052~3.827 | 5.293* | Not significant |

All data reflect n = 25 for YA, and n = 22 for OA independent participants. *p < 0.05, **p < 0.01, *** p< 0.001.

Table S6: Mediation analysis using interregional informational connectivity as a mediator, calculated by averaging interregional informational connectivity across unique edges from the Old-Adult atlas (including ERC-CA1, PRC-CA1, SUB-CA1, and CA2/3/DG-CA1), with the neural remapping index of CA1 as an independent variable and spatial distance discrimination performance as a dependent variable (results from the Old-Adult atlas).

| Covariate | c<br>Total effect | a | b | a*b<br>Mediation effect | a*b<br>(95% BootCI) | c'<br>Direct effect | Conclusion |
|---|---|---|---|---|---|---|---|
| CA1 Volume and Gender | 12.258** | -4.781* | -0.584* | 2.793 | -1.235~9.149 | 9.465* | Partial mediation |
| CA1 Volume, Gender and Age | 8.985** | -3.109 | -0.204 | 0.633 | -1.185~4.756 | 8.352* | Not significant |

All data reflect n = 25 for YA, and n = 22 for OA independent participants. *p < 0.05, **p < 0.01.

Table S7: Mediation analysis using interregional informational connectivity as a mediator, calculated by averaging interregional informational connectivity across common edges for both the Young-Adult and the Old-Adult atlas (including ERC-CA1, PRC-CA1, SUB-CA1 and CA2/3/DG-CA1), with the neural remapping index of CA1 as an independent variable and spatial distance discrimination performance as a dependent variable (results from the Young-Adult atlas).

| Covariate | c<br>Total effect | a | b | a*b<br>Mediation effect | a*b<br>(95% BootCI) | c'<br>Direct effect | Conclusion |
|---|---|---|---|---|---|---|---|
| CA1 Volume and Gender | 9.314** | -3.358** | -0.874* | 2.934 | 0.857~6.091 | 6.380* | Partial mediation |
| CA1 Volume, Gender and Age | 6.592** | -2.659* | -0.514 | 1.368 | -0.118~4.225 | 5.224* | Not significant |

All data reflect n = 25 for YA, and n = 22 for OA independent participants. *p < 0.05, **p < 0.01.

Table S8: Mediation analysis using interregional informational connectivity as a mediator, calculated by averaging interregional informational connectivity across common edges for both the Young-Adult and the Old-Adult atlas (including ERC-CA1, PRC-CA1, SUB-CA1 and CA2/3/DG-CA1), with the neural remapping index of CA1 as an independent variable and spatial distance discrimination performance as a dependent variable (results from the Old-Adult atlas).

| Covariate | c<br>Total effect | a | b | a*b<br>Mediation effect | a*b<br>(95% BootCI) | c'<br>Direct effect | Conclusion |
|---|---|---|---|---|---|---|---|
| CA1 Volume and Gender | 12.258** | -4.781* | -0.584* | 2.793 | -1.235~9.149 | 9.465* | Partial mediation |
| CA1 Volume, Gender and Age | 8.985** | -3.109 | -0.204 | 0.633 | -1.185~4.756 | 8.352* | Not significant |

All data reflect n = 25 for YA, and n = 22 for OA independent participants. *p < 0.05, **p < 0.01.

Table S9: Mediation analysis using interregional informational connectivity of PRC-ERC as a mediator, neural remapping index of CA1 as an independent variable and spatial distance discrimination performance as a dependent variable (results from the Young-Adult atlas).

| Covariate | c<br>Total effect | a | b | a*b<br>Mediation effect | a*b<br>(95% BootCI) | c'<br>Direct effect | Conclusion |
|---|---|---|---|---|---|---|---|
| CA1 Volume and Gender | 9.314** | -6.859* | -0.235* | 1.613 | -0.115~4.523 | 7.701** | Partial mediation |
| CA1 Volume, Gender and Age | 6.592** | -4.485 | -0.1 | 0.448 | -0.653~2.626 | 6.144* | Not significant |

All data reflect n = 25 for YA, and n = 22 for OA independent participants. *$p < 0.05$, **$p < 0.01$.

Table S10: Mediation analysis using interregional informational connectivity of PRC-ERC as a mediator, neural remapping index of CA1 as an independent variable and spatial distance discrimination performance as a dependent variable (results from the Old-Adult atlas).

| Covariate | c<br>Total effect | a | b | a*b<br>Mediation effect | a*b<br>(95% BootCI) | c'<br>Direct effect | Conclusion |
|---|---|---|---|---|---|---|---|
| CA1 Volume and Gender | 12.258** | -11.283** | -0.377** | 4.259 | 0.399~11.089 | 7.999* | Partial mediation |
| CA1 Volume, Gender and Age | 8.985** | -8.689* | -0.164 | 1.429 | -1.180~6.169 | 7.556* | Not significant |

All data reflect n = 25 for YA, and n = 22 for OA independent participants. *p < 0.05, **p < 0.01.

Table S11: Mediation analysis using interregional informational connectivity for the same spatial environment as a mediator, calculated by averaging interregional informational connectivity across unique edges from the Young-Adult atlas (including PHC-CA1, ERC-CA1 and SUB-CA1), with standard deviation of within-city MPS in CA1 as an independent variable and spatial distance discrimination performance as a dependent variable (results from the Young-Adult atlas).

| Covariant | c<br>Total effect | a | b | a*b<br>Mediation effect | a*b<br>(95% BootCI) | c'<br>Direct effect | Conclusion |
|---|---|---|---|---|---|---|---|
| CA1 Volume and Gender | -11.023* | 4.779* | -0.822* | -3.93 | -10.187~0.1771 | -7.092 | Full mediation |
| CA1 Volume, Gender and Age | -2.77 | 2.587 | -0.457 | -1.182 | -5.778~0.953 | -1.588 | Not significant |

All data reflect n = 25 for YA, and n = 22 for OA independent participants. *$p < 0.05$.

Table S12: Mediation analysis using interregional informational connectivity for the same environment as a mediator, calculated by averaging interregional informational connectivity across common edges for both the Young-Adult and the Old-Adult atlases (including ERC-CA1, SUB-CA1 and CA2/3/DG-CA1), with standard deviation of within-city MPS in CA1 as an independent variable and spatial distance discrimination performance as a dependent variable (results from the Young-Adult atlas).

| Covariant | c<br>Total effect | a | b | a*b<br>Mediation effect | a*b<br>(95% BootCI) | c'<br>Direct effect | Conclusion |
|---|---|---|---|---|---|---|---|
| CA1 Volume and Gender | -11.023* | 4.284* | -0.934* | -4 | -9.025~-0.287 | -7.023 | full mediation |
| CA1 Volume, Gender and Age | -2.77 | 2.636 | -0.552 | -1.455 | -5.701~0.824 | -1.315 | Not significant |

All data reflect n = 25 for YA, and n = 22 for OA independent participants. *$p < 0.05$.

Table S13: Mediation analysis using interregional informational connectivity for the same environment as a mediator, calculated by averaging interregional informational connectivity across unique edges from the Old-Adult atlas (including ERC-CA1, PRC-CA1 and SUB-CA1), with standard deviation of within-city MPS in CA1 as an independent variable and spatial distance discrimination performance as a dependent variable (results from the Old-Adult atlas).

| Covariant | c<br>Total effect | a | b | a*b<br>Mediation effect | a*b<br>(95% BootCI) | c'<br>Direct effect | Conclusion |
|---|---|---|---|---|---|---|---|
| CA1 Volume and Gender | -5.095 | 6.484* | -0.578* | -3.749 | -10.987~0.029 | -1.345 | Full mediation |
| CA1 Volume, Gender and Age | -2.801 | 5.267* | 0.034 | 0.18 | -3.559~4.371 | -2.981 | Not significant |

All data reflect n = 25 for YA, and n = 22 for OA independent participants. *$p < 0.05$.

Table S14: Mediation analysis using interregional informational connectivity for the same environment as a mediator, calculated by averaging interregional informational connectivity across common edges for both the Young-Adults and the Old-Adult atlases (including ERC-CA1, SUB-CA1 and CA2/3/DG-CA1), with standard deviation of within-city MPS in CA1 as an independent variable and spatial distance discrimination performance as a dependent variable (results from the Old-Adult atlas).

| Covariant | c<br>Total effect | a | b | a*b<br>Mediation effect | a*b<br>(95% BootCI) | c'<br>Direct effect | Conclusion |
|---|---|---|---|---|---|---|---|
| CA1 Volume and Gender | -5.095 | 6.394** | -0.591~ | -3.777 | -12.283~0.037 | -1.317 | full mediation |
| CA1 Volume, Gender and Age | -2.801 | 5.418** | 0.038 | 0.208 | -4.104~5.127 | -3.009 | Not significant |

All data reflect n = 25 for YA, and n = 22 for OA independent participants. **p < 0.01.

Table S15 Mediation analysis using dimensionality of CA1 as a mediator, neural remapping index of CA1 as an independent variable and spatial distance discrimination as a dependent variable (results from the Young-Adult atlas).

| Covariate | c<br>Total effect | a | b | a*b<br>Mediation effect | a*b<br>(95% BootCI) | c'<br>Direct effect | Conclusion |
|---|---|---|---|---|---|---|---|
| CA1 Volume and Gender | 1068.316** | 0.008** | 8.191 | 0.089 | 5.098~12.104 | 1.123 | Full mediation |
| CA1 Volume, Gender and Age | 6.592** | 863.080*** | 0.006*** | 5.577 | 2.776~8.882 | 1.015 | Full mediation |

All data reflect n = 25 for YA, and n = 22 for OA independent participants. **$p < 0.01$, ***$p < 0.001$.

Table S16 Mediation analysis using dimensionality of CA1 as a mediator, neural remapping index of CA1 as an independent variable and spatial distance discrimination as a dependent variable (results from the Old-Adult atlas).

| Covariate | c<br>Total effect | a | b | a*b<br>Mediation effect | a*b<br>(95% BootCI) | c'<br>Direct effect | Conclusion |
|---|---|---|---|---|---|---|---|
| CA1 Volume and Gender | 12.258** | 1388.225** | 0.008** | 11.006 | 5.734~18.747 | 1.252 | Full mediation |
| CA1 Volume, Gender and Age | 8.985*** | 1157.294*** | 0.006*** | 7.482 | 3.677~13.209 | 1.503 | Full mediation |

All data reflect n = 25 for YA, and n = 22 for OA independent participants. **$p < 0.01$, ***$p < 0.001$.

Table S17. Behavioral information during learning session prior MRI scanner for each participant.

| ParticipantID | Group | City1_runs | City2_runs | City1 ACC (%) | City2 ACC (%) | Overall ACC |
|---|---|---|---|---|---|---|
| 102 | YA | 3 | 3 | 1 | 1 | 1 |
| 103 | YA | 3 | 3 | 0.6 | 0.8 | 0.7 |
| 105 | YA | 3 | 3 | 1 | 1 | 1 |
| 106 | YA | 3 | 3 | 1 | 1 | 1 |
| 107 | YA | 3 | 3 | 0.9 | 0.8 | 0.85 |
| 108 | YA | 3 | 3 | 1 | 1 | 1 |
| 109 | YA | 3 | 3 | 0.9 | 0.8 | 0.85 |
| 110 | YA | 3 | 3 | 0.8 | 0.9 | 0.85 |
| 111 | YA | 3 | 3 | 1 | 1 | 1 |
| 112 | YA | 4 | 4 | 0.9 | 0.9 | 0.9 |
| 113 | YA | 3 | 3 | 0.9 | 0.8 | 0.85 |
| 114 | YA | 3 | 3 | 0.9 | 0.9 | 0.9 |
| 115 | YA | 3 | 3 | 0.9 | 0.8 | 0.85 |
| 116 | YA | 3 | 3 | 0.7 | 1 | 0.85 |
| 117 | YA | 3 | 3 | 0.8 | 0.9 | 0.85 |
| 118 | YA | 3 | 3 | 0.9 | 0.8 | 0.85 |
| 119 | YA | 3 | 3 | 0.9 | 0.9 | 0.9 |
| 120 | YA | 3 | 3 | 1 | 0.9 | 0.95 |
| 121 | YA | 3 | 3 | 1 | 1 | 1 |
| 123 | YA | 3 | 3 | 1 | 0.9 | 0.95 |
| 126 | YA | 3 | 3 | 1 | 0.7 | 0.85 |
| 127 | YA | 3 | 3 | 0.9 | 1 | 0.95 |
| 128 | YA | 3 | 3 | 1 | 1 | 1 |
| 129 | YA | 3 | 3 | 1 | 1 | 1 |
| 131 | YA | 3 | 3 | 0.8 | 0.7 | 0.75 |
| $Mean_{YA}$ | YA | 3.04 | 3.04 | 0.912 | 0.9 | 9.06 |
| 202 | OA | 6 | 6 | 0.7 | 0.7 | 0.7 |
| 203 | OA | 3 | 3 | 0.7 | 0.7 | 0.7 |
| 204 | OA | 3 | 3 | 0.8 | 0.9 | 0.85 |
| 205 | OA | 4 | 4 | 0.7 | 0.7 | 0.7 |
| 206 | OA | 7 | 6 | 0.6 | 0.7 | 0.65 |
| 207 | OA | 6 | 6 | 0.9 | 0.8 | 0.85 |
| 208 | OA | 3 | 3 | 0.7 | 0.8 | 0.75 |
| 210 | OA | 8 | 9 | 0.9 | 0.7 | 0.8 |
| 211 | OA | 3 | 4 | 0.7 | 1 | 0.85 |
| 212 | OA | 3 | 3 | 0.8 | 0.7 | 0.75 |
| 215 | OA | 5 | 3 | 0.8 | 0.9 | 0.85 |
| 216 | OA | 3 | 3 | 1 | 0.8 | 0.9 |
| 217 | OA | 7 | 6 | 0.9 | 0.8 | 0.85 |
| 218 | OA | 4 | 4 | 0.9 | 0.9 | 0.9 |
| 219 | OA | 3 | 3 | 0.7 | 0.8 | 0.75 |

| | | | | | | |
|---|---|---|---|---|---|---|
| 220 | OA | 3 | 3 | 0.9 | 0.9 | 0.9 |
| 221 | OA | 5 | 3 | 0.7 | 0.6 | 0.65 |
| 222 | OA | 3 | 3 | 0.9 | 0.8 | 0.85 |
| 223 | OA | 3 | 3 | 0.7 | 0.8 | 0.75 |
| 224 | OA | 3 | 3 | 1 | 1 | 1 |
| 226 | OA | 3 | 3 | 0.7 | 0.9 | 0.8 |
| 227 | OA | 3 | 3 | 0.7 | 0.9 | 0.8 |
| $Mean_{OA}$ | OA | 4.136 | 3.954 | 0.791 | 0.809 | 0.8 |

City1_runs: the number of runs for City 1 completed by each participant during the learning session to reach the criteria. City2_runs: the number of runs for City 2 completed by each participant during the learning session to reach the criteria. The participantIDs highlighted in red represent older adults who did not meet a more stringent behavioral criterion (i.e., accuracy of 75%) during the training stage. YA: young adults, OA: old adults.

Table S18. The count of observations/trials that involved in different analyses.

| Group | Mixed effect model/ Interregional informational Connectivity analysis | | PCA |
|---|---|---|---|
| | within-city trial pairs | between-city trial pairs | trials |
| Young adults | 1706.7 ± 480.1 | 2609.5 ± 731.8 | 101.2 ± 15.5 |
| Old adults | 986.1 ± 632.6 | 1493.9 ± 962.2 | 72.8 ± 28.2 |
| Overall | 1369.4 ± 659.6 | 2087.3 ± 1009.3 | 87.9 ± 26.3 |
